# Improving tolerance to fluctuating light through adaptive laboratory evolution in the cyanobacterium Synechocystis

**DOI:** 10.1101/2025.07.20.665829

**Authors:** Theo Figueroa-Gonzalez, Eslam M. Abdel-Salam, Weiyang Chen, Milena Zhivkovikj, Marcel Dann, Dario Leister

**Author notes:** **Correspondence** and requests for materials should be addressed to Dario Leister.

## Abstract

Fluctuating light (FL) poses a serious challenge to photosynthetic organisms like cyanobacteria, disrupting carbon assimilation and damaging photosystems. While key components of cyanobacterial FL tolerance have been identified, their genetic enhancement remains unexplored. We applied adaptive laboratory evolution to *Synechocystis* PCC 6803 under two complex FL regimes, including one lethal to the starter strain (LT), to generate FL-adaptive alleles. Our analysis revealed 44 fully segregated novel mutations in 24 monoclonal evolved strains, 28 of which affected proteins or structural RNAs. We focused on three mutations for further study. Mutations in Pam68, involved in photosystem II (PSII) assembly, and Sll0518 were present in all evolved strains, indicating early emergence. These mutations increased tolerance to non-lethal FL conditions when introduced into LT, with the Pam68 mutation possibly protecting PSI by increasing the proportion of less active PSII monomers. A gain-of-function mutation in RpaB, regulator of phycobilisome association B, was found in three strains tolerant to lethal FL. When introduced into LT, this mutation significantly increased tolerance to both lethal FL and high light conditions, associated with downregulation of photosystem accumulation and light harvesting. As RpaB has plant homologs, this finding could potentially be used to improve agricultural productivity under variable light conditions.

## INTRODUCTION

Cyanobacteria are unique prokaryotes capable of oxygenic photosynthesis. Photosynthesis in cyanobacteria involves photosystem I (PSI) and II (PSII) complexes, which capture light energy and drive water-splitting and reductive equivalent production. This process creates a proton gradient across thylakoid membranes, enabling ATP synthesis and CO_2_ fixation via the Calvin-Benson-Bassham cycle^1,2^. While light is essential for photosynthesis, excessive or fluctuating light can be detrimental, causing photoinhibition^3,4^. Cyanobacteria often encounter high light (HL) and fluctuating light (FL) conditions in their aquatic and terrestrial habitats, with FL photon flux densities potentially reaching extreme levels^5,6,7^. Extensive research has been conducted on cyanobacterial HL tolerance mechanisms, including gene expression^8,9,10,11,12,13^, proteomic^14,15,16^ and metabolic^17,18^ responses, PSI structural adaptations^6^, non-photochemical quenching^19^, and effects of mutations in individual proteins^20,21^. However, FL tolerance and acclimation in cyanobacteria remain less explored.

Studies on FL-exposed cultures of the model cyanobacterial species *Synechococcus elongatus* PCC 7942 and *Synechocystis* sp. PCC 6803 (hereafter “Synechocystis”) have revealed the importance of inorganic carbon availability and alternative electron pathways for FL tolerance^22,23,24^. Flavodiiron (Flv) proteins mediate an alternative electron pathway, enhancing FL tolerance by reducing O_2_ and alleviating PSI acceptor site limitation without reactive oxygen species production^23,25,26^. Transcriptomic and proteomic responses suggest nitrogen assimilation acts as an electron sink, protecting the photosynthetic electron transport chain during FL^27,28^. Thylakoid-localized respiratory terminal oxidase activity may alleviate sudden photosynthetic electron availability during FL, limiting photoinhibition^29^ Additionally, Fluctuating-light acclimation protein 1 (FLAP1) increases FL tolerance by negatively regulating light-induced proton extrusion, preventing cytosolic over-alkalization during dark-to-light transitions^30^.

Although more FL tolerance components likely exist, no genetic enhancement of cyanobacterial FL tolerance has been reported, hindering the development of suitable production strains for FL-prone photobioreactors^31^. Similarly, improving flowering plant FL tolerance through genetic engineering has seen limited success, with a few exceptions in tobacco and soybean^32^. Further increases in acclimation potential may require an evolutionary approach entailing the identification of new FL tolerance factors and the evolution of advantageous alleles^33^.

Previous adaptive laboratory evolution (ALE) studies on Synechocystis for increased HL tolerance have demonstrated the general accessibility of photosynthetic robustness to evolutionary improvement^34,35,36^. Therefore, we applied ALE to evolve novel alleles conferring FL tolerance in Synechocystis, resulting in the identification of distinct candidate mechanisms for tolerance to different types of light fluctuations. Mutation of the cyanobacteria-specific protein Sll0518 and of Pam68, which is involved in the early assembly of PSII and is conserved between cyanobacteria and flowering plants, conferred tolerance to moderate FL conditions. The drastically increased tolerance to both FL and HL conditions conferred by a mutation in RpaB, regulator of phycobilisome association B, which has homologs in eukaryotic algae and flowering plants, holds promise for potential application in plants to enhance FL tolerance and photosynthetic efficiency.

## RESULTS

### Generation of FL-tolerant batch cultures through adaptive evolution

We adapted Synechocystis strains to tolerate fluctuating light (FL) using adaptive laboratory evolution (ALE), building on previous methods for generating high light (HL) tolerance^33^. Based on previous findings^33^, we relied on the natural mutation rate of Synechocystis without using external mutagens.

Two experimental protocols were designed to progressively increase FL intensity. These protocols modified an existing FL regimen for *Arabidopsis thaliana*^37^, which alternated between high and low light phases. The first protocol, “FL0”, maintained the original 1-min HL / 5-min LL rhythm but increased light intensity. Starting with 700 μmol photons m^-2^ s^-1^ for HL (HL_700_) and 50 μmol photons m^-2^ s^-1^ for LL (LL_50_), the intensity of the HL and LL phases were gradually altered to 1200 (HL_1200_) and 12 μmol photons m^-2^ s^-1^ (LL_12_), respectively (**Supplementary Table 1, Fig. 1a**). While continuous HL_1200_ was lethal to non-adapted strains, the FL0 final conditions allowed for recovery and growth during the LL phase (**Fig. 1b**).

**Fig. 1.**
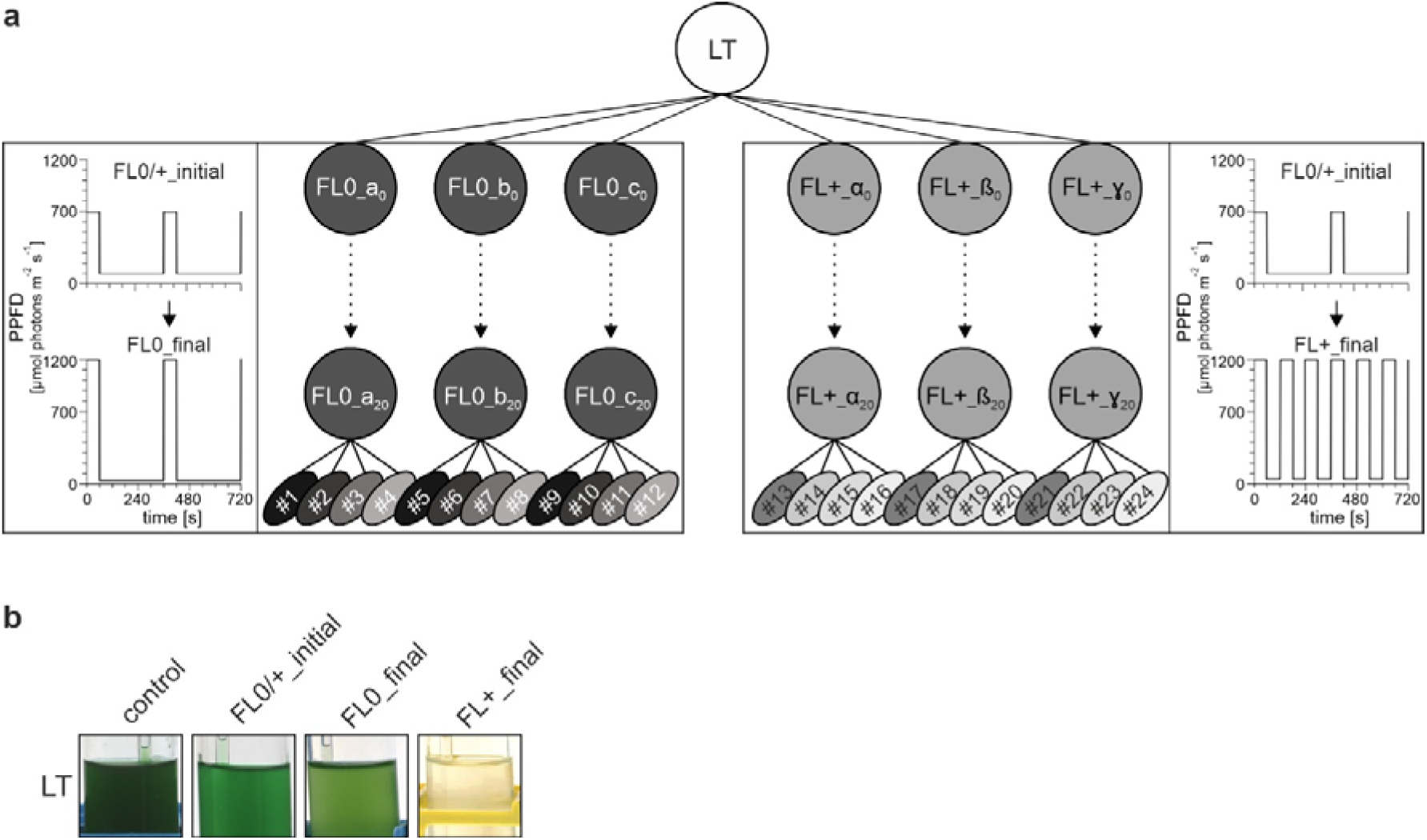
Experimental design of the two FL-ALE experiments in Synechocystis. **a** ALE scheme. Batch cultures are shown as circles, propagation events as dots, and monoclonal cultures as ovals. A glucose-tolerant, non-motile laboratory type (LT) of *Synechocystis* sp. PCC 6803 was used to initiate six independent batch cultures, three for each protocol (FL0_a_0_, FL0_b_0_, FL0_c_0_ and FL+_a_0_, FL+_b_0_, aFL+_c_0_). The FL0 protocol initially subjected cultures to cycles of 1 min high light (HL) (700 μmol photons m^-2^ s^-1^, HL_700_) followed by 5 min low light (LL) (50 μmol photons m^-2^ s^-1^, LL_50_). Over 20 cycles, the light intensity amplitude was gradually increased, culminating in 12 final cycles of 1 min HL_1200_ and 5 min LL_12_. The FL+ protocol initially mirrored FL0, but progressively reduced the LL phase duration to 1 min while increasing the light intensity amplitude. The final 12 cycles consisted of 1 min HL_1200_ alternating with 1 min LL_12_. The illumination protocols are detailed as part of the flow chart. After 20 months and 20 propagation cycles, the resulting cultures (FL0_a_20_, FL0_b_20_, FL0_c_20_ and FL+_α_20_, FL+_β_20_, FL+_γ_20_) were characterized. Four monoclonal isolates from each of the two sets of triplicate batch culture (#1 to #24, shown as ovals in different shades of grey to represent variability) were subjected to whole-genome sequencing and further analysis. **b** Image of the original LT culture grown under various illumination conditions: constant LL_50_(control), initial fluctuating light for both protocols (FL0/+ initial, 1 min HL_700_ and 5 min LL_50_), final FL0 protocol (FL0_final, 1 min HL_1200_ and 5 min LL_12_) and the lethal lethal FL+ conditions (FL+_final, 1 min HL_1200_ and 1 min LL_12_). Cultures were maintained at 23 °C with atmospheric aeration (100-150 mL air per min).

The second protocol, “FL+”, began with the same initial conditions as FL0 but progressively shortened the LL phase to 1 min (**Supplementary Table 1, Fig. 1a**). The final cycles alternated between 1 min at LL_12_ and 1 min at HL_1200_. These conditions proved lethal for non-adapted strains (**Fig. 1b**). Both FL0 and FL+ protocols involved 20 selective cultivation cycles on triplicate batch cultures over 20 months.

After adaptive evolution, the cultures showed phenotypic variations within and between triplicates (**Fig 2a-d**). Under their respective final FL conditions, FL+ strains exhibited higher growth rates and increased cell density compared to FL0 cultures, although with lower chlorophyll content. This suggests that FL+ adapted strains not only tolerated the lethal light regime but also utilized the greater available light energy more effectively than FL0 strains.

**Fig. 2.**
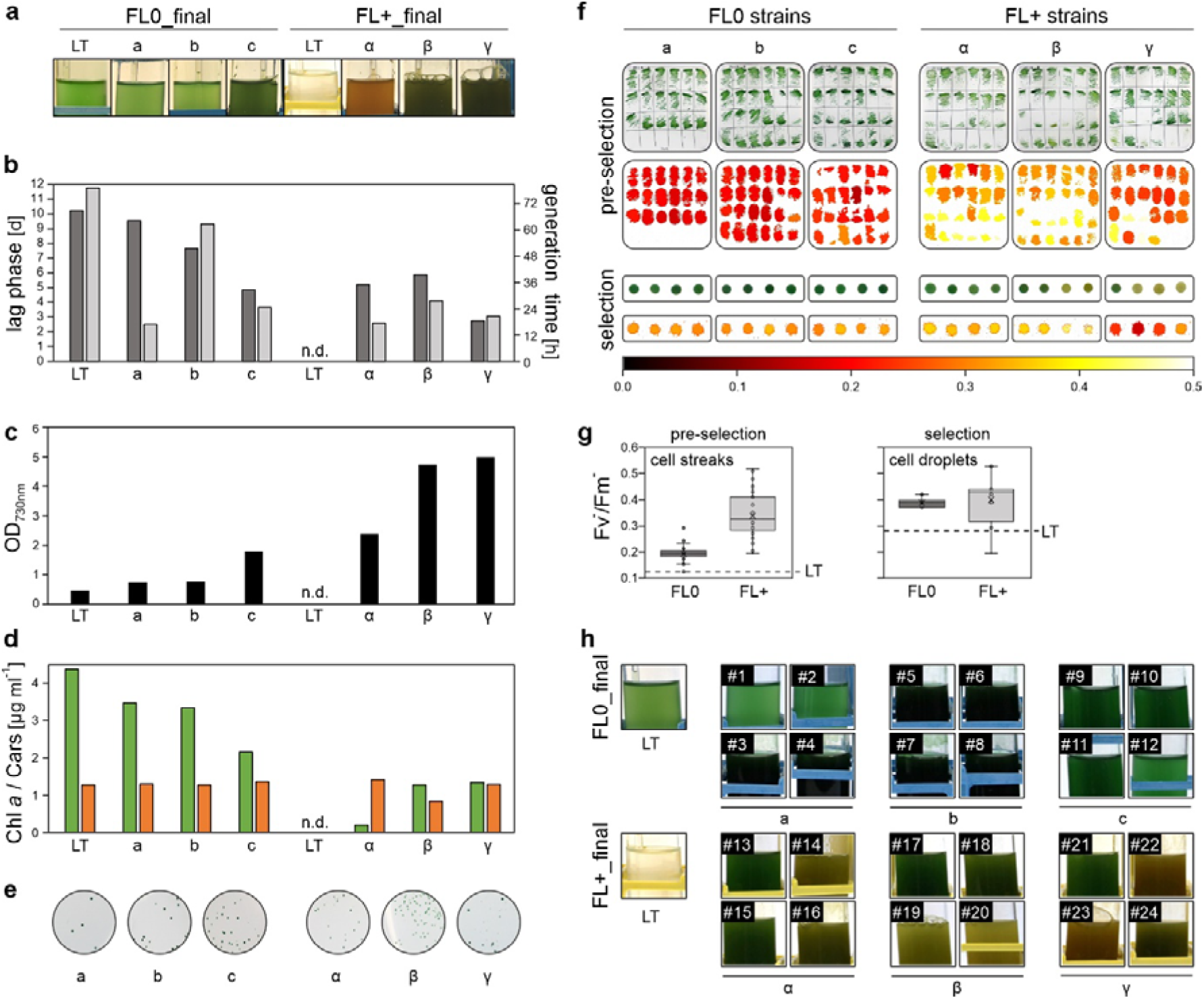
Characterisation of final FL-ALE batch cultures and derived monoclonal strains. **a-d** The six independently adapted final batch cultures (FL0: a, b, c; FL+: α, β, γ) were cultivated for 14 days under their respective final FL regimes. Visual appearance (**a**), growth kinetics (lag phase and generation time) (**b**), cell density (**c**), and photosynthetic pigment content (chlorophyll (Chl) a and carotenoids) (**c**) were quantified. **e** Monoclonal isolation was performed by serial dilution (1:10^6^-10^7^) of the adapted cultures, followed by plating on BG11 agar and incubation under constant LL_20_. **f** Individual clones were screened in two stages. Pre-selection: Clones were subcultured on agar plates for 7 days under constant LL_20_ at 23 °C. Colony morphology and chlorophyll fluorescence (Fv^-^/Fm^-^) were assessed. Selection: Four clones spanning the Fv^-^/Fm^-^ range were chosen from each pre-selected set. These were grown in liquid culture, normalized to OD_730nm_=10, and 10 µL aliquots were spotted on agar plates for further growth and Fv^-^/Fm^-^ analysis. **g** Statistical analysis of Fv^-^/Fm^-^ values of pre-selected (n = 66 for FL0, 72 for FL+) and selected (n = 12 each) clones was performed. Dotted lines indicate Fv^-^/Fm^-^ values for LT control. Note that the difference in the two LT Fv^-^/Fm^-^ values between the two measurements is, at least in part, due to the different cell densities of the cell streaks and normalized droplets^70^. **h** The 24 selected clones (12 FL0, #1 to #12; 12 FL+, #13 to #24) were cultivated for 14 days under their respective selective final light regimes, with LT controls for comparison. Box plots show individual data points, median (horizontal lines), mean (crosses), interquartile range (box), and 1.5× interquartile range (whiskers).

### Isolation and characterization of FL-tolerant monoclonal strains

To isolate individual clones, the FL0 and FL+ batch cultures were diluted, plated on solid media, and incubated under constant LL_20_ conditions (**Fig. 2e**). After picking and transferring individual clones to new plates, they were allowed to grow for 7 days before imaging and measuring the ’apparent’ quantum yield of PSII, Fv^-^/Fm^-^^38^ (**Fig. 2f**). It is important to note that measuring and analyzing Fv/Fm in cyanobacteria can be problematic due to the contribution of phycobilisomes to basal fluorescence and interference from respiratory components^38,39^. The FL+ clones exhibited greater heterogeneity in Fv^-^/Fm^-^ values compared to FL0 clones (**Fig. 2f, g**).

Four clones from each batch were selected to represent the quantiles of Fv^-^/Fm^-^, sampling a wide range of phenotypic diversity. Liquid cultures of these clones were normalized and applied to solid media for further fluorescence data collection (**Fig. 2f**). FL0 isolates showed a homogeneous phenotype with dark green color and Fv^-^/Fm^-^ of 0.39 ± 0.02. In contrast, FL+ clones displayed clear heterogeneity, with colors ranging from cyan to ochre and Fv-/Fm- values between 0.19 and 0.53 (0.40 ± 0.10) (**Fig. 2f, g**).

The 24 monoclonal cultures were grown for two weeks under their respective selective regimes. Most FL0 isolates grew denser than the LT control under FL0 final conditions, while all FL+ clones survived and accumulated high cell densities under the final FL+ protocol, which was lethal to the non-adapted LT (**Fig. 2h**).

In summary, the monoclonal strains derived from FL0 and FL+ batch cultures demonstrated improved growth rates compared to LT under their respective selective conditions. The FL+ monoclonal strains exhibited marked phenotypic heterogeneity in terms of coloration and Fv^-^/Fm^-^ values, highlighting the diverse adaptations that occurred during the selection process.

### Mutations in FL-tolerant monoclonal strains

We conducted a comprehensive whole-genome analysis of the 12 FL0 and 12 FL+ monoclonal strains, using the LT strain from which these adapted strains were derived and the original motile *Synechocystis* PCC 6803 strain (designated “WT”) as controls. Genome sequence analysis yielded a mutation matrix revealing 412 mutations (234 in FL0 and 269 in FL+) absent in both LT and WT strains (**Supplementary** Fig. 1a).

The majority of these mutations (349 total, 201 in FL0 and 223 in FL+) were located within coding regions of the genome. Almost all (342 total, 198 in FL0 and 218 in FL+) were single nucleotide polymorphisms (SNPs), while seven (3 in FL0 and 5 in FL+) were insertions or deletions (InDels). Among the coding region SNPs, 277 (157 in FL0 and 182 in FL+) resulted in non-synonymous exchanges, modifying the amino acid sequence of 89 proteins (53 and 56 in FL0 and FL+, respectively) with known function and affecting an additional 188 proteins (104 and 126 in FL0 and FL+, respectively) with unknown functions. A comparative analysis among the 24 strains showed considerable variability in the ratios of non-synonymous to synonymous mutations **(Supplementary** Fig. 1b).

The distribution of allele frequencies across the two FL-ALE experiments was extremely polarized: almost two-thirds of the mutations were classified as ’low-frequency’ occurring in ≤ 10% of the reads, while about one-quarter of the alleles were fully segregated, exhibiting a frequency of 100%, indicating their presence in virtually every read from the respective samples (**Supplementary** Fig. 1c). In contrast to the allele frequencies reported for the subset of the HL-ALE experiment which did not include external mutagens^49^, the high proportion of low-frequency alleles appears to be characteristic of our FL-ALE experiment (**Supplementary** Fig. 1c).

### FL adaptive haplotype

Genomic analysis identified 101 fully segregated mutations that were present in the evolved strains, LT, or WT, but absent from the published Synechocystis reference genome. Phylogenetic reconstruction based on these mutations showed a clear distinction between adapted strains and controls **(Fig. 3**). FL0 strains shared 39 fully segregated mutations, including 6 novel alleles (absent in LT/WT controls), and exhibited shorter genetic distances among themselves compared to FL+ clones. FL+ strains displayed increased phenotypic variability, as reflected by larger genetic distances.

**Fig. 3.**
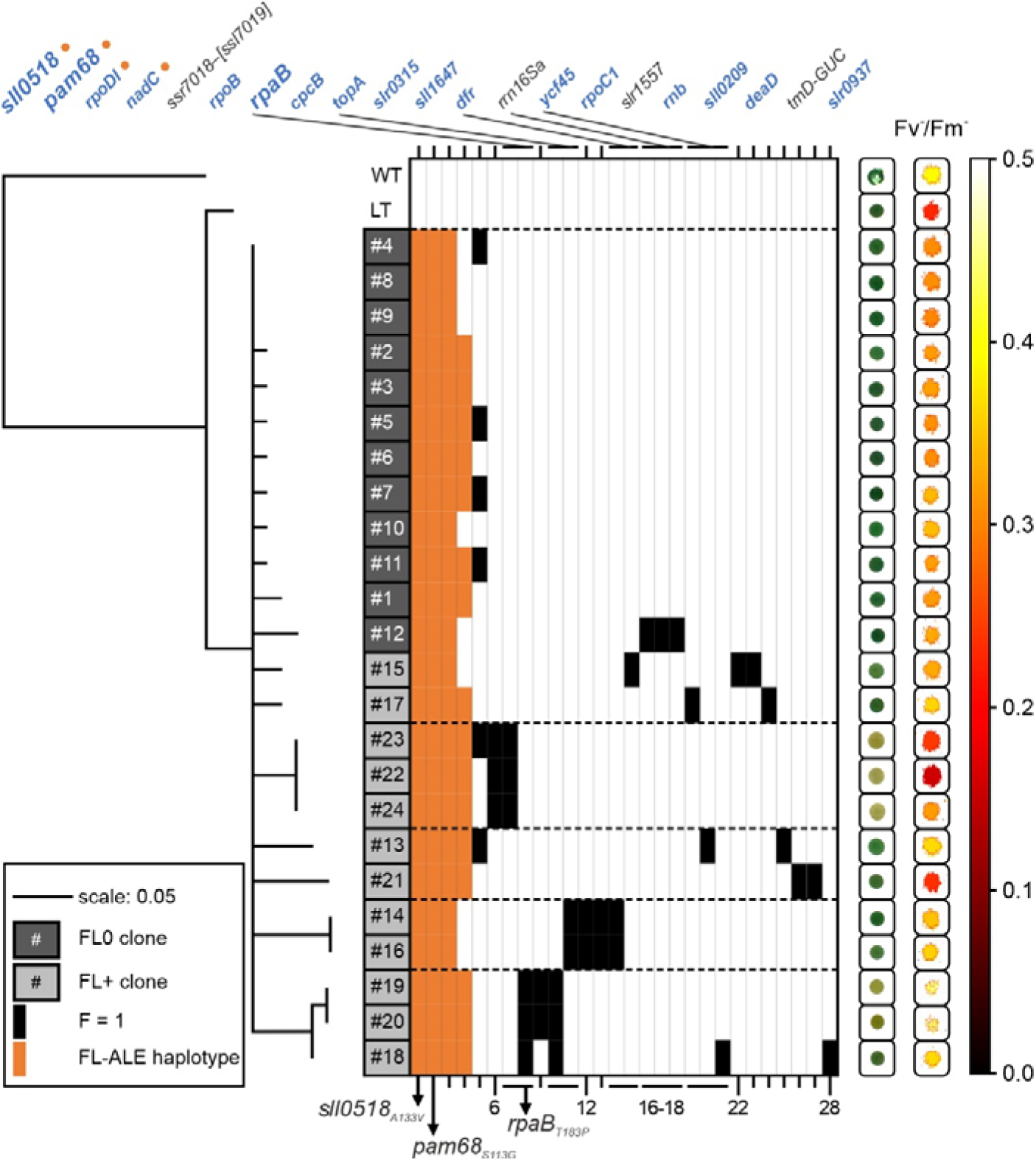
Phylogenetic and mutational analysis of the monoclonal FL-ALE strains. The left panel displays a maximum likelihood phylogram illustrating the genetic relationship among FL0-ALE (dark grey boxes) and FL+-ALE (light grey boxes) monoclonal strains. The strain numbers (#) correspond to the ones in **Fig. 2h**. The phylogram includes the original motile, glucose-sensitive *Synechocystis* sp. PCC 6803 isolate (WT) and the laboratory type (LT, non-motile, glucose-tolerant) as reference points. Genetic distances are represented by horizontal solid lines, with a scale of 0.05 substitutions per site. The phylogenetic analysis is based on 101 fully segregated alleles (allele frequency = 1), with only branches showing ≥80% bootstrap support displayed. The evolved strains are grouped into five distinct clades, demarcated by horizontal dashed lines. The central panel depicts a map of 28 mutations absent in WT and LT strains. This includes 24 protein-altering mutations (20 non-synonymous base substitutions and 4 indels) and 4 mutations affecting rRNAs or tRNAs (refer to **Table 1** for details). Mutations present at 100% frequency in at least one monoclonal strain are represented by black or orange rectangles, with orange indicating alleles common to ≥16 of the 24 strains. Loci are arranged left to right based on mutation frequency across the 24 strains, except for mutations affecting the same locus, which are grouped together. Genes affected by non-synonymous SNPs are highlighted in blue. Gene identifiers separated by “/” denote deletions in intergenic regions, while “–” indicates deletions affecting coding regions (affected genes in brackets). Three key alleles - *sll0518_A133V_*, *pam68_S113G_*, and *rpaB_T183P_* - are emphasized in bold and marked by arrow-heads, respectively. The right panel presents photographic images of representative cell droplets for each monoclonal strain, accompanied by their respective Fv^-^/Fm^-^ values, providing a visual and quantitative phenotypic characterization of the evolved strains.

**Table 1.** Fully segregated, protein-affecting mutations identified in FL-ALE strains. The table catalogs non-synonymous single nucleotide polymorphisms (SNPs) and deletions within coding regions that are absent in both the laboratory type (LT) and wild-type (WT) strains, and have achieved 100% allele frequency in at least one of the 24 monoclonal strains. Columns are organized as follows: “#”: Corresponds to the mutation numbering in **Fig. 3**. “Locus”: Gene identifier; “–” denotes deletions within coding regions, with affected genes in brackets “[]”. “Mutation”: Specifies the nature and position of SNPs or indels, using the notation “x/y nt” where x is the affected position in a sequence of total length y. “No.”: Indicates the number of monoclonal strains harboring the specific allele. “ALE”: Specifies the ALE protocol(s) in which the allele was detected (0 for FL0, + for FL+, 0+ for both). “Function”: Assigns Gene Ontology (GO) terms - M (metabolic processes), P (photosynthesis), Tc (transcription), Tl (translation), and(unknown function). Alleles associated with the FL-tolerant haplotype (present in ≥65% of monoclonal strains) are highlighted in orange. Alleles reconstituted and assessed for FL tolerance in this study are highlighted in bold font.

| # | Locus | Annotation | Mutation | No. | ALE | Function |
| --- | --- | --- | --- | --- | --- | --- |
| 1 | *sll0518* | unknown protein | A133V (GCC→GTC) | 24 | 0+ | ? |
| 2 | *pam68* (*sll0933*) | PAM68 protein | S113G (AGC→GGC) | 24 | 0+ | P |
| 3 | *rpoDI* (*slr0653*) | RNA polymerase $\sigma$ factor | R96L (CGT→CTT) | 24 | 0+ | Tc |
| 4 | *nadC* (*slr0936*) | nicotinatenucleotide pyrophosphorylase | Q183H (CAA→CAT) | 16 | 0+ | M |
| 5 | *ssr7018*–[*ssl7019*] | hypothetical proteins | Δ935 bp | 6 | 0+ | ? |
| 6 | *rpoB* (*sll1787*) | RNA polymerase beta subunit | T791N (ACC→AAC) | 3 | + | Tc |
| 7 | *rpaB* (*slr0947*) | **OmpR subfamily** | **T183P (ACC→CCC)** | **3** | + | **Tc** |
| 8 |  |  | D194G (GAC→GGC) | 3 | + |  |
| 9 | *cpcB* (*sll1577*) | phycocyanin b subunit | S40P (TCT→CCT) | 2 | + | P |
| 10 | *topA* (*slr2058*) | DNA topoisomerase I | T504A (ACC→GCC) | 3 | + | Tc |
| 11 |  |  | E149D (GAA→GAT) | 2 | + |  |
| 12 | *slr0315* | hypothetical protein | V62L (GTT→CTT) | 2 | + | ? |
| 13 | *sll1647* | hypothetical protein | W97R (TGG→AGG) | 2 | + | ? |
| 14 | *dfr* (*sll0698*) | drug sensory protein A | T393A (ACC→GCC) | 2 | + | M |
| 15 |  |  | F605L (TTC→TTG) | 1 | + |  |
| 16 | *rrn16Sa* | 16S ribosomal RNA | noncoding (254/1489 nt) | 1 | 0 | Tl |
| 17 |  |  | noncoding (247/1489 nt) | 1 | 0 |  |
| 18 |  |  | noncoding (243/1489 nt) | 1 | 0 |  |
| 19 | *ycf45* (*slr0692*) | hypothetical protein | Δ13 bp, coding (331-343/1770 nt) | 1 | + | ? |
| 20 |  |  | E533G (GAA→GGA) | 1 | + |  |
| 21 |  |  | +T, coding (59/1770 nt) | 1 | + |  |
| 22 | *rpoC1 (slr1265)* | RNA polymerase<br>$\gamma$ subunit | R221L (CGG→CTG) | 1 | + | Tc |
| 23 | *slr1557* | unknown protein | Δ8 bp, coding (573-580/1110 nt) | 1 | + | ? |
| 24 | *rnb (sll1290)* | ribonuclease II | V500M (GTG→ATG) | 1 | + | Tc |
| 25 | *sll0209* | hypothetical protein | Q290E (CAA→GAA) | 1 | + | ? |
| 26 | *deaD (slr0083)* | ATP dependent RNA<br>helicase; DeaD | R399L (CGG→CTG) | 1 | + | Tc |
| 27 | *trnD-GUC* | tRNA for aspartate | A→G, noncoding (8/74 nt) | 1 | + | Tl |
| 28 | *slr0937* | unknown protein | G292S (GGT→AGT) | 1 | + | ? |

Of the 101 fully segregated mutations, 44 were novel (not present in LT or WT), including 16 mutations in non-coding regions, 24 protein-altering mutations, and four mutations in structural RNAs. Of these 28 fully segregated mutations, affecting proteins or structural RNAs, five were common to both FL0 and FL+ strains, three were specific to FL0 strains, and 20 were specific to FL+ strains (**Table 1**, **Fig. 3**). Three mutations (in *rpoDI*, *sll0518*, *pam68*) occurred in all 24 evolved strains, and five loci harbored multiple mutations. Functionally, the mutated genes were involved in various processes, including transcription (6), translation (2), metabolism (2), and photosynthesis (2) (**Table 1**).

Three mutations were selected for further analysis: *sll0518_A133V_*, a missense mutation in a gene encoding a cyanobacteria-specific protein of unknown function, *pam68_S113G_*, a missense mutation in *pam68*/*sll0933*, encoding a factor involved in the early assembly of PSII^40^, and *rpaB_T183P_,* a missense mutation in the *rpaB*/*slr0947* gene, encoding the regulator of phycobilisome association B, the inactivation of which reduces the efficiency of energy transfer from phycobilisomes to PSII^41^. The *sll0518_A133V_*and *pam68_S113G_* mutations were common to all 24 evolved strains, suggesting that they arose early in the adaptive process. The RpaB_T183P_ mutation was exclusive to three FL+ strains, and its importance in conferring FL+ tolerance is further supported by the independent occurrence of another mutation of RpaB (RpaB_D194G_) in three other FL+ strains (**Table 1**, **Fig. 3**).

In conclusion, the genomic analysis revealed a higher frequency of fully segregated, protein-altering mutations in FL+ strains compared to FL0 strains. A substantial subset of these mutations was exclusive to FL+ strains, indicating more extensive genetic adaptation to the stringent FL+ conditions. Conversely, fewer fully segregated mutations were specific to FL0 strains. The shared mutations between FL+ and FL0 strains, particularly those present in all evolved strains, suggest their importance in conferring improved tolerance to the initial FL conditions common to both protocols.

### Recapitulating FL tolerance in LT cells and interplay of FL and HL tolerance

To assess the individual contributions of mutations identified through FL0 and FL+ adaptive ALE to FL tolerance, three specific mutations (*sll0518_A133V_*, *pam68S_113G_*, and *rpaB_T183P_*) were introduced into the LT strain. The growth of these mutant strains was then evaluated under final FL0 and final FL+ conditions. The *sll0518_A133V_* and *pam68_S113G_* mutants showed significantly enhanced FL0 tolerance compared to the LT control, with cell density increases of 107% and 131%, respectively (**Fig. 4**). However, these strains failed to grow productively under final FL+ conditions. In contrast, *rpaB_T183P_* displayed growth comparable to LT under FL0 conditions but outperformed all other strains under final FL+ conditions (**Fig. 4)**.

**Fig. 4.**
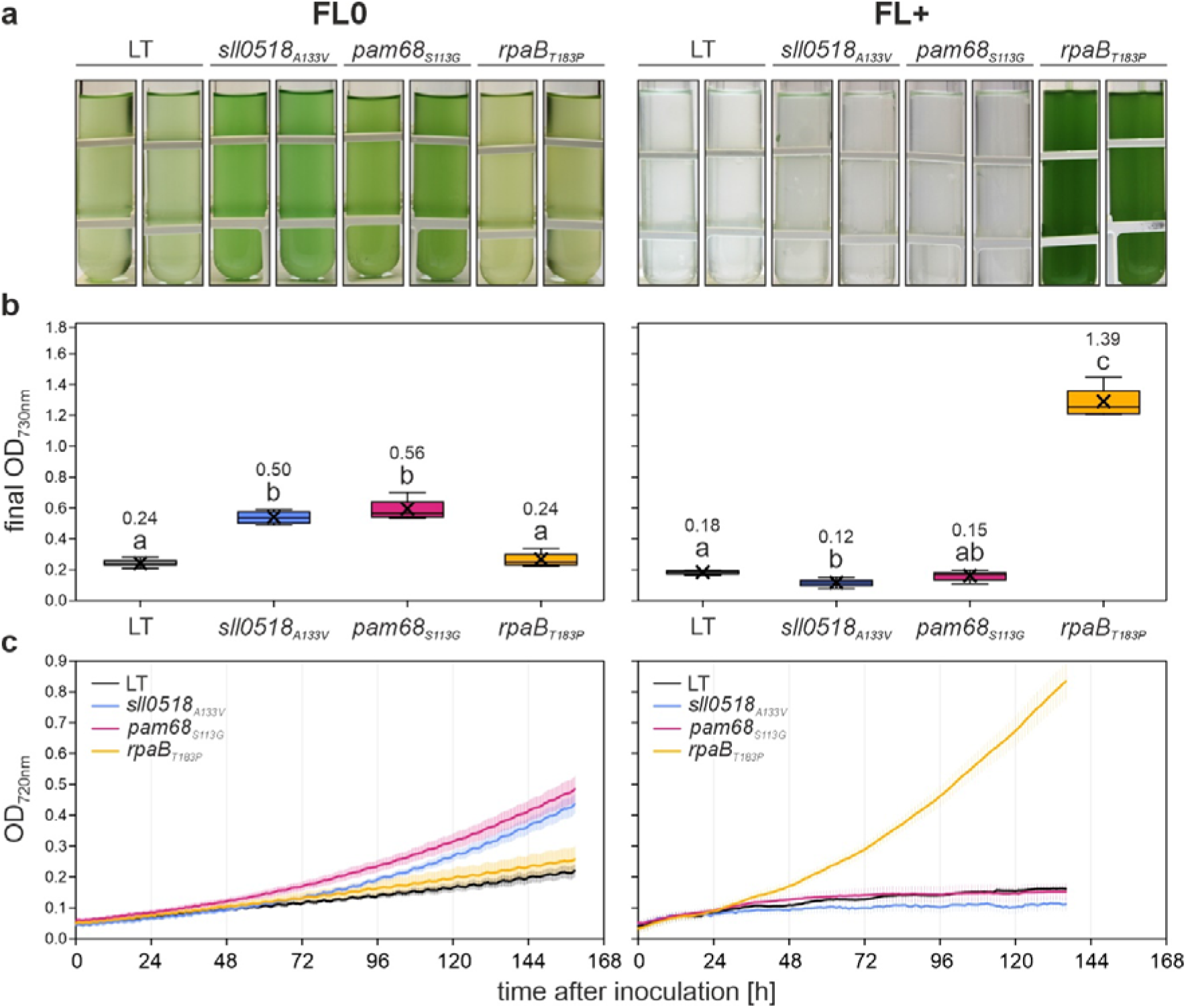
Growth characteristics of *sll0518_A133V_*, *pam68_S113G_*, *rpaB_T183P_* and LT strains under final FL0 and FL+ conditions. a Visual representation of liquid cultures for the four strains cultivated in multi-cultivators under final FL0 (left) or FL+ (right) conditions at 23 °C with 100 mL min^-1^ aeration, photographed seven days post-inoculation. **b** Quantitative analysis of cell density (OD_730nm_) for the four strains grown under conditions described in A. Box plots are presented, with lowercase letters denoting statistically significant differences (p ≤ 0.05) as determined by post-hoc Bonferroni-Holm simultaneous comparison of all measurements (n = 8) following significant between-group differences detected by one-factorial ANOVA. **c** Growth kinetics of the four strains under final FL0 and FL+ conditions, monitored automatically by multi-cultivators measuring OD_720nm_. Error bars represent the standard deviation (n = 4, except for *sll0518_A133V_* and *pam68_S113G_* under FL+, where n = 8). Statistical data in panel b are visualized using box plots, showing individual data points, median (horizontal lines), mean (crosses), interquartile range (box), and 1.5× interquartile range (whiskers).

To explore whether these mutations also affect growth under constant light intensities, the mutant strains and LT control were cultured under constant low light (LL_12_) and constant high light (HL_1200_) conditions. Under LL_12_, *sll0518_A133V_*exhibited slightly increased growth, *rpaB_T183P_* showed slightly decreased growth compared to LT, while *pam68_S113G_* grew similarly to LT (**Supplementary** Fig. 2). Under HL_1200_ conditions, which are lethal for non-evolved LT cells, only *rpaB_T183P_* demonstrated productive growth, while LT, *sll0518_A133V_*, and *pam68_S113G_* failed to grow (**Supplementary** Fig. 2).

To investigate the potential correlation between tolerance to final FL+ conditions and constant HL (HL_1200_) conditions, we examined whether mutations conferring HL tolerance could also impart resistance to FL+ conditions. This hypothesis was based on the observation that the *rpaB_T183P_* strain exhibited tolerance to both final FL+ and constant HL_1200_ conditions. To test this hypothesis, we evaluated the growth of three previously characterized strains: HL1, HL2, and HL1/2^35^. Contrary to our hypothesis, none of the HL-tolerant strains exhibited growth under FL+ conditions (**Supplementary** Fig. 3). This observation suggests that the mechanisms underlying tolerance to constant HL and fluctuating HL conditions may be distinct, and that adaptations to one condition do not necessarily confer tolerance to the other.

In summary, the results indicate that two mutations (*sll0518_A133V_*and *pam68_S113G_*) identified in all strains adapted to final FL0 or FL+ conditions confer enhanced tolerance to FL0 when individually introduced into the LT strain. However, these mutations alone do not provide tolerance to FL+ conditions. Conversely, a mutation specific to FL+ adaptation (*rpaB_T183P_*) imparts tolerance to both FL+ and constant HL conditions, even in the absence of the four common mutations found in the FL-adapted haplotypes. This suggests that the FL-adapted haplotype mutations are not prerequisite for FL+ or HL tolerance. The results also reveal that mutations conferring HL tolerance do not necessarily provide tolerance to FL+ conditions, which consist of alternating 1-min pulses of HL intensity and LL intensity, respectively. This observation, coupled with the fact that LT strains can grow under FL0 conditions (with a short HL phase followed by a longer LL phase) but not under continuous HL_1200_, indicates that extended LL phases enable cells to withstand otherwise lethal light intensities. In contrast, brief LL phases appear to exacerbate the detrimental effects of the HL phase.

### The Pam68_S113G_ mutation enhances photosynthetic protein accumulation under LL and stabilizes monomeric PSII under FL

The *pam68_S113G_* mutation, involving a serine-to-glycine substitution at position 113 of the Sll0933/Pam68 protein, demonstrated enhanced tolerance to final FL0 conditions compared to the LT progenitor strain. To investigate the functional implications of this mutation, we created a Pam68 overexpression strain (*pam68oe*) controlled by the *psbA2* promoter and compared its growth characteristics with an insertional knockout mutant for *pam68* (*ins0933*)^40^ under both LL_12_ and final FL0 conditions (**Fig. 5**). Under FL0, *ins0933* achieved a final OD_730nm_ 32% higher than LT, while *pam68_S113G_* reached 92% higher. The *pam68oe* strain attained a final OD_730nm_ 67% higher than LT on average, slightly less than *pam68_S113G_* but not significantly different from *ins0933* and *pam68_S113G_*. Under LL_12_ conditions, no significant growth differences were observed among the strains **(Fig. 5**). These findings suggest that the *pam68_S113G_* mutation enhances FL tolerance similarly to WT Pam68 overexpression.

**Fig. 5.**
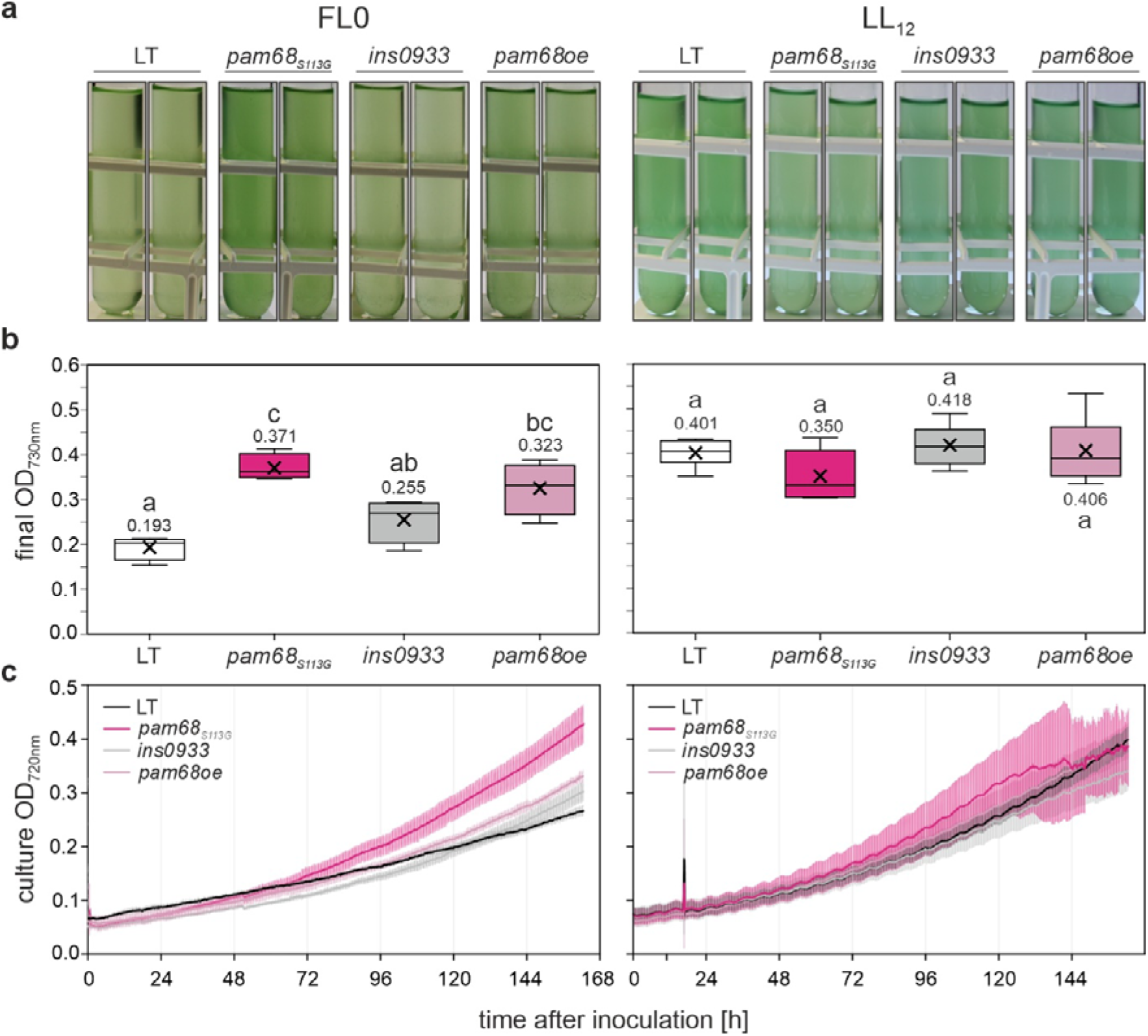
Growth characteristics of the *ins0933*, oe*Pam68*, *pam68_S113G_*, and LT strains under final FL0 and constant LL_12_ conditions. **a** Visual representation of liquid cultures for the four strains cultivated in multi-cultivators under final FL0 and constant LL_12_ conditions at 23 °C with 100 mL min^-1^ aeration, photographed seven days post-inoculation. **b** Quantitative analysis of cell density (OD_730nm_) for the four strains grown under conditions as in A. Box plots are presented, with lowercase letters denoting statistically significant differences (p ≤ 0.05) as determined by *post-hoc* Bonferroni-Holm simultaneous comparison of all measurements (*n* = 4 for FL0 condition and *n* = 6 for LL_12_ condition, respectively) following significant between-group differences detected by one-factorial ANOVA. **c** Growth kinetics of the four strains under final FL+ conditions, monitored automatically by multi-cultivators measuring OD_720nm_. Error bars represent the standard deviation (*n* = 4 for FL0 condition and *n* = 6 for LL_12_ condition, respectively). Statistical data in panel b are presented as box plots, showing individual data points, median (horizontal lines), mean (crosses), interquartile range (box), and 1.5× interquartile range (whiskers).

Given the non-essential nature of Sll0933/Pam68 as a PSII assembly factor in Synechocystis^40^, its interactions with YCF48, CP43, and CP47, and its potential role in CP47 and CP43 assembly and PSII core complex formation^42^, we examined thylakoid protein levels. LT cultures and two independent *pam68_S113G_* mutant strains were grown under continuous LL_50_ and final FL0 conditions. Under LL_50_, *pam68_S113G_* cultures showed a 73% increase in endpoint OD_730nm_ compared to LT (2.78 ± 0.09 vs. 1.61 ± 0.05), while under FL0, *pam68_S113G_* exhibited an even stronger increase (0.59 ± 0.07 vs. 0.24 ± 0.02) as already seen in Fig. 5 (**Fig. 6a**). Immunoblot analysis assessed the accumulation of Pam68 and representative thylakoid proteins (D1 and PsaC), while allophycocyanin (APC) and phycocyanin (PC) levels were quantified via in-gel fluorescence (**Fig. 6b**). Under LL_50_, all four proteins showed significant increases in *pam68_S113G_* compared to LT. Under FL0 conditions, *pam68_S113G_* cells displayed reduced Pam68 protein levels compared to LT (-24 %), while other protein levels remained largely unaltered. These results suggest increased accumulation of photosynthetic complexes under constant LL_50_ in *pam68_S113G_* mutant strains, with no such effect under FL0 conditions (**Fig. 6c**). This indicates *pam68_S113G_* as a likely gain-of-function mutation, given that its beneficial effect on growth under FL0 conditions exceeds Pam68 overexpression (**Fig. 5**) despite lowered Pam68 protein levels.

**Fig. 6.**
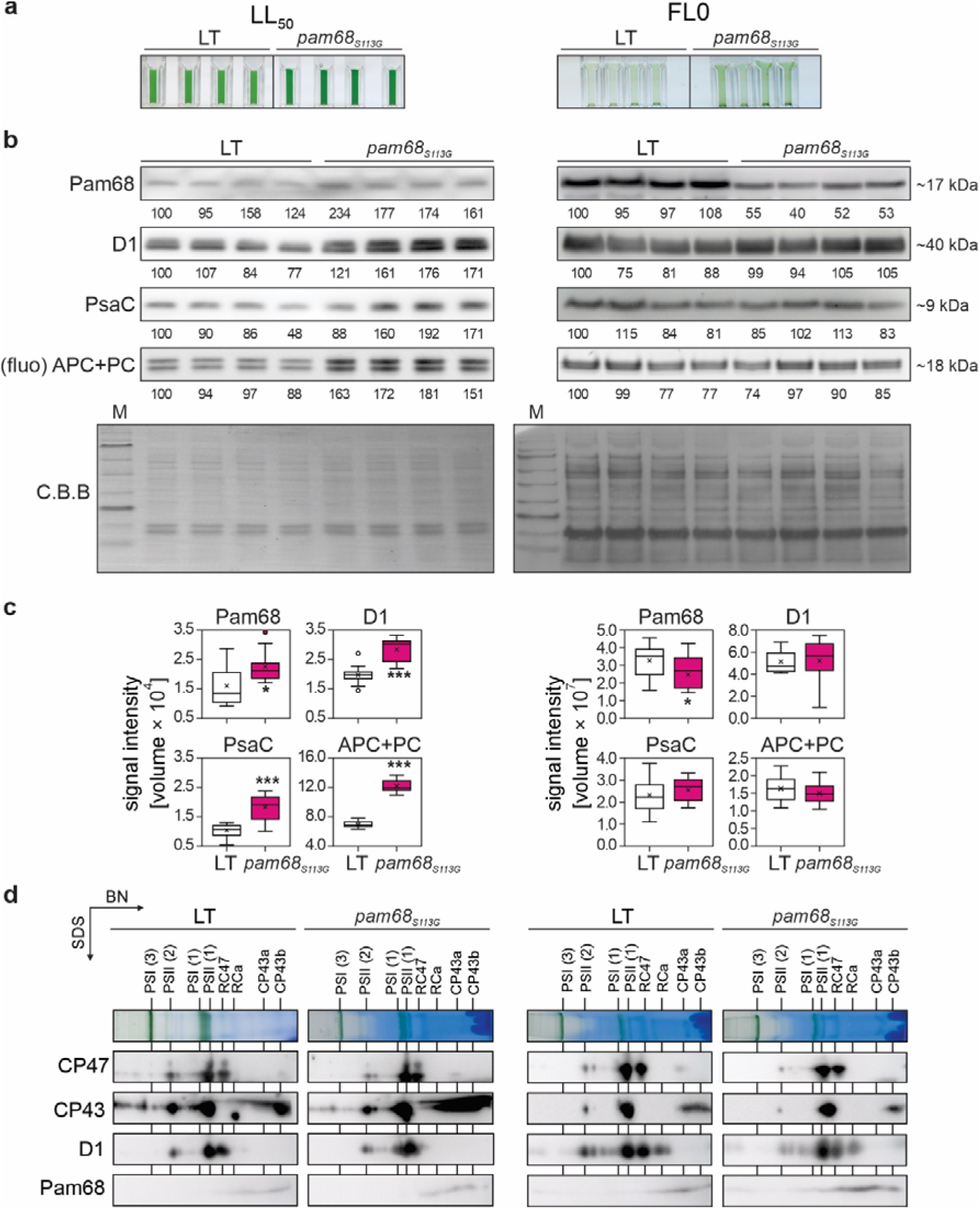
The *pam68_S113G_* mutation enhances PSII accumulation at LL. a Visual representation of liquid cultures for the LT and *pam68_S113G_* strains cultivated under constant LL_50_ and final FL0 conditions at 23 °C with 100 mL min^-1^ aeration, photographed seven days post-inoculation. **b** Immunoblot analysis. Whole-cell protein extracts were obtained from the cultures shown in panel a. Aliquots containing ∼10 µg of total protein content, as determined by Bradford assay, were subjected to SDS-PAGE separation. Chromophore fluorescence from phycobilisome marker proteins (allophycocyanin and phycocyanin, APC+PC) was visualized directly from the gels. Proteins were electro-transferred onto PVDF membranes. Specific antibodies were employed for immunodetection of Pam68, D1, and PsaC. A Coomassie Brilliant Blue (C.B.B) stain of the PVDF membrane served as a loading control. Relative quantification of immunoblot signals was performed, with values normalized to the first lane of each blot. Comprehensive data analysis is presented in panel C. Molecular weight markers are indicated. **c** Quantification of signals obtained for Pam68, D1, PsaC, and APC+PC (n = 12, except for APC+PC, where n = 24). Data are presented as box plots, showing individual data points, median (horizontal lines), mean (crosses), interquartile range (box), and 1.5× interquartile range (whiskers). Statistical significance was determined using Student’s t-Test, with p ≤ 0.05 (*), p ≤ 0.01 (**), and p ≤ 0.001 (***). **d** Two-dimensional PAGE (BN/SDS-PAGE) analysis of thylakoid proteins. Thylakoid preparations (equivalent to ∼5 µg chlorophyll) were solubilized and separated in two dimensions. Proteins were electro-transferred onto PVDF membranes for immunodetection using specific antibodies against D1, CP43, CP47, and Pam68. PSII assembly complexes are annotated as follows: monomer/dimer/trimer (1/2/3), PSII core complex lacking CP43 (RC47), reaction center complex lacking PSII core antennae modules CP43 and CP47 (RCa), and CP43 module a/b (CP43a/b)^42^.

To investigate the role of Pam68_S113G_ in PSII level increase under LL_50_ and potential effects under FL0, we analyzed PSII complex assembly using two-dimensional PAGE (**Fig. 6d**). Overall, no pronounced differences in mature complexes or assembly intermediates of PSII were observed. CP43-containing PSII assembly intermediates of lower molecular weight (CP43a and b) were increased in *pam68_S113G_* under LL_50_ conditions, but not under FL0. The *pam68_S113G_* mutant showed a clear decrease in dimeric PSII abundance compared to monomeric PSII under FL0 conditions. Pam68 was predominantly found in the low molecular weight fraction, with no observable depletion of thylakoid-associated Pam68 protein levels in *pam68_S113G_* relative to LT under FL0, despite the lowered total levels of *pam68_S113G_* (see **Fig. 6b**).

These observations suggest that an altered Pam68 functionality in *pam68_S113G_* mutants may promote physical interaction with PSII assembly intermediates and stabilize monomeric PSII at the expense of PSII dimer formation. Since Synechocystis PSII monomers exhibit lower activity compared to PSII dimers^43^, the enhanced FL0 tolerance observed in the *pam68_S113G_* strain could partly result from indirect PSI protection due to reduced PSII electron pressure.

### The *rpaB_T183P_* mutation is associated with downregulation of light harvesting

RpaB/Slr0947, an essential response regulator in *Synechocystis* sp. PCC 6803, plays a crucial role in regulating energy transfer from phycobilisomes to photosystems^44,45,46^. This regulatory function suggests that reduced light harvesting capacity may contribute to the enhanced tolerance of *rpaB_T183P_* cultures to HL and FL conditions that are lethal to the starter strain. The *rpaB_T183P_*mutant strain, characterized by a threonine-to-proline substitution at residue 183 of the RpaB/Slr0947 protein, showed increased tolerance to final FL+ conditions that are lethal for the LT progenitor strain. To investigate the functional implications of this mutation, an RpaB overexpression strain (*rpaBoe*) was generated and compared with a previously characterized RpaB knock-down mutant (*rpaBkd*)^46^ under constant HL_1200_ and final FL+ conditions. Notably, only the *rpaB_T183P_* mutant demonstrated productive growth under both conditions, while LT, *rpaBkd*, and *rpaBoe* strains failed to grow (**Fig. 7**). Further investigation of the RpaB_T183P_ mutation revealed that under LL_50_ conditions, LT and *rpaB_T183P_* had similar endpoint OD_730nm_ values (1.61 ± 0.05 and 1.64 ± 0.09, **Fig. 8a**). However, under HL_700_ conditions, *rpaB_T183P_* showed markedly higher values (1.78 ± 0.36 vs. 4.19 ± 0.60), and under FL+ conditions, only *rpaB_T183P_* grew (1.29 ± 0.10) while LT failed (**Fig. 8a**).

**Fig. 7.**
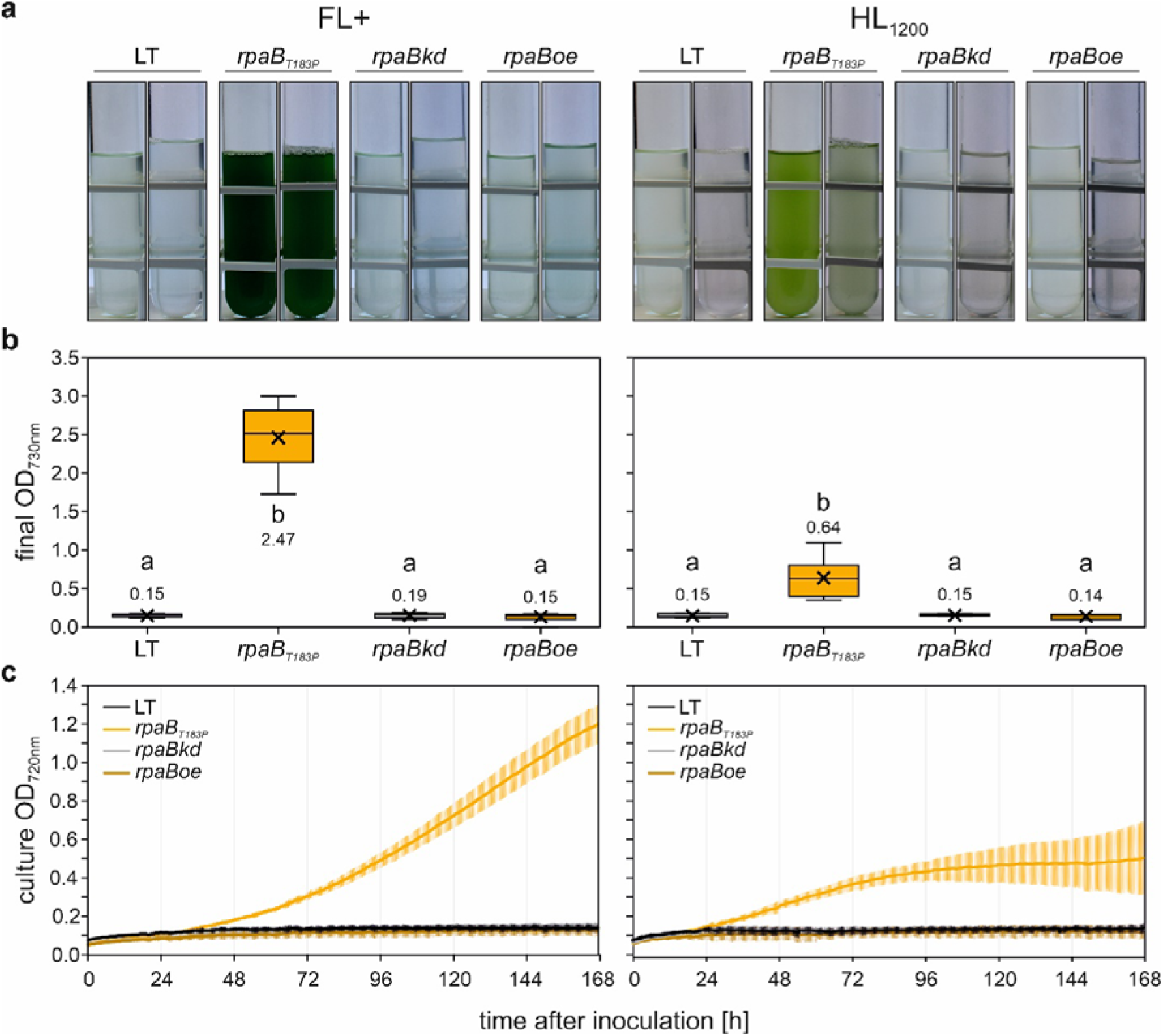
Growth characteristics of the *rpaBkd*, *rpaBoe*, *rpaB_T183P_*, and LT strains under final FL+ and constant HL_1200_ conditions. a Visual representation of liquid cultures for the four strains cultivated in multi-cultivators under final FL+ and constant HL_1200_ conditions at 23 °C with 100 mL min^-1^ aeration, photographed seven days post-inoculation. **b** Quantitative analysis of cell density (OD_730nm_) for the four strains grown under conditions as in A. Box plots are presented, with lowercase letters denoting statistically significant differences (p ≤ 0.05) as determined by post-hoc Bonferroni-Holm simultaneous comparison of all measurements following significant between-group differences detected by one-factorial ANOVA. **c** Growth kinetics of the four strains under final FL+ conditions, monitored automatically by multi-cultivators measuring OD_720nm_. Error bars represent the standard deviations. FL+ data shown in panels b and c represents *n* = 5/8/5/4 biological replicates of LT/*rpaB_T183P_*/*rpaB_kd_*/*rpaB_oe_*, respectively. HL_1200_ data shown in panels B and C represents n = 4/6/4/3 biological replicates of LT/*rpaB_T183P_*/*rpaB_kd_*/*rpaB_oe_*, respectively. Statistical data in panel B are presented as box plots, showing individual data points, median (horizontal lines), mean (crosses), interquartile range (box), and 1.5× interquartile range (whiskers).

**Fig. 8.**
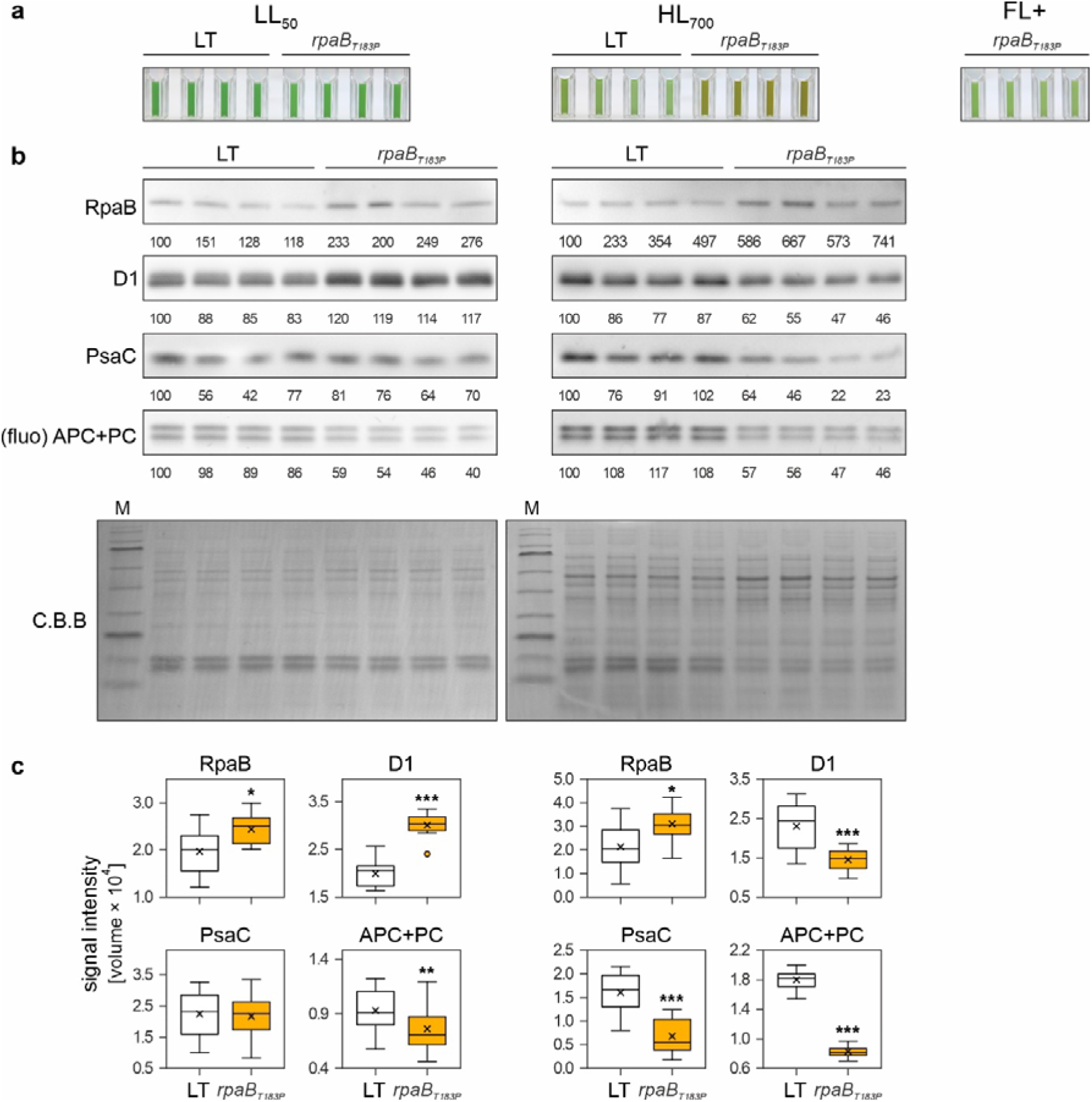
The *rpaB_T183P_* mutation is associated with downregulation of light harvesting. a Visual representation of liquid cultures for the LT and *rpaB_T183P_* strains cultivated under constant LL_50_, constant HL_700_ and final FL+ (*rpaB_T183P_* only) at 23 °C with 100 mL min^-1^ aeration, photographed seven days post-inoculation. **b** Immunoblot analysis. Whole-cell protein extracts were obtained from the cultures shown in panel a. Aliquots containing ∼10 µg of total protein content, as determined by Bradford assay, were subjected to SDS-PAGE separation. Chromophore fluorescence from phycobilisome marker proteins (allophycocyanin and phycocyanin, APC+PC) was visualized directly from the gels. Proteins were electro-transferred onto PVDF membranes. Specific antibodies were employed for immunodetection of Pam68, D1, and PsaC. A Coomassie Brilliant Blue (C.B.B) stain of the PVDF membrane served as a loading control. Relative quantification of immunoblot signals was performed, with values normalized to the first lane of each blot. Comprehensive data analysis is presented in panel c. Molecular weight markers are indicated. **c** Quantification of signals obtained for Pam68, D1, PsaC, and APC+PC (n = 12, except for APC+PC, where n = 24). Data are presented as box plots, showing individual data points, median (horizontal lines), mean (crosses), interquartile range (box), and 1.5× interquartile range (whiskers). Statistical significance was determined using Student’s t-Test, with p ≤ 0.05 (*), p ≤ 0.01 (**), and p ≤ 0.001 (***).

Immunoblot analysis and in-gel fluorescence quantification showed slightly increased RpaB levels in *rpaB_T183P_* under LL_50_ and HL_700_ compared to LT. At LL_50_, D1 levels increased while PsaC levels decreased in *rpaB_T183P_* (**Fig. 8b,c)**. Under HL_700_, all thylakoid proteins (D1, PsaC, and APC+PC) were significantly downregulated in *rpaB_T183P_*, indicating a general downregulation of photosynthesis.

In vivo light energy harvesting capacity was investigated through absorption spectra of intact cells (**Fig. 9a**). The ratio between PC and Chl a absorption maxima was slightly reduced in *rpaB_T183P_* under LL_50_ conditions and more pronounced reduced under HL_700_ conditions, indicating downregulation of light harvesting. Fluorescence emission spectra at low temperature (77 K), determined by exciting with phycobilisome-specific wavelengths (λ_max_ = 600 nm) and calculating the total signal as the sum of fluorescence intensities between 626 to 780 nm, showed reduced phycobilisome-mediated energy allocation in *rpaB_T183P_* cultures (**Fig. 9b**, likely due to reduced phycobilisome antenna size and consistent with the reduced APC+PC levels (**Fig. 8c**).

**Fig. 9.**
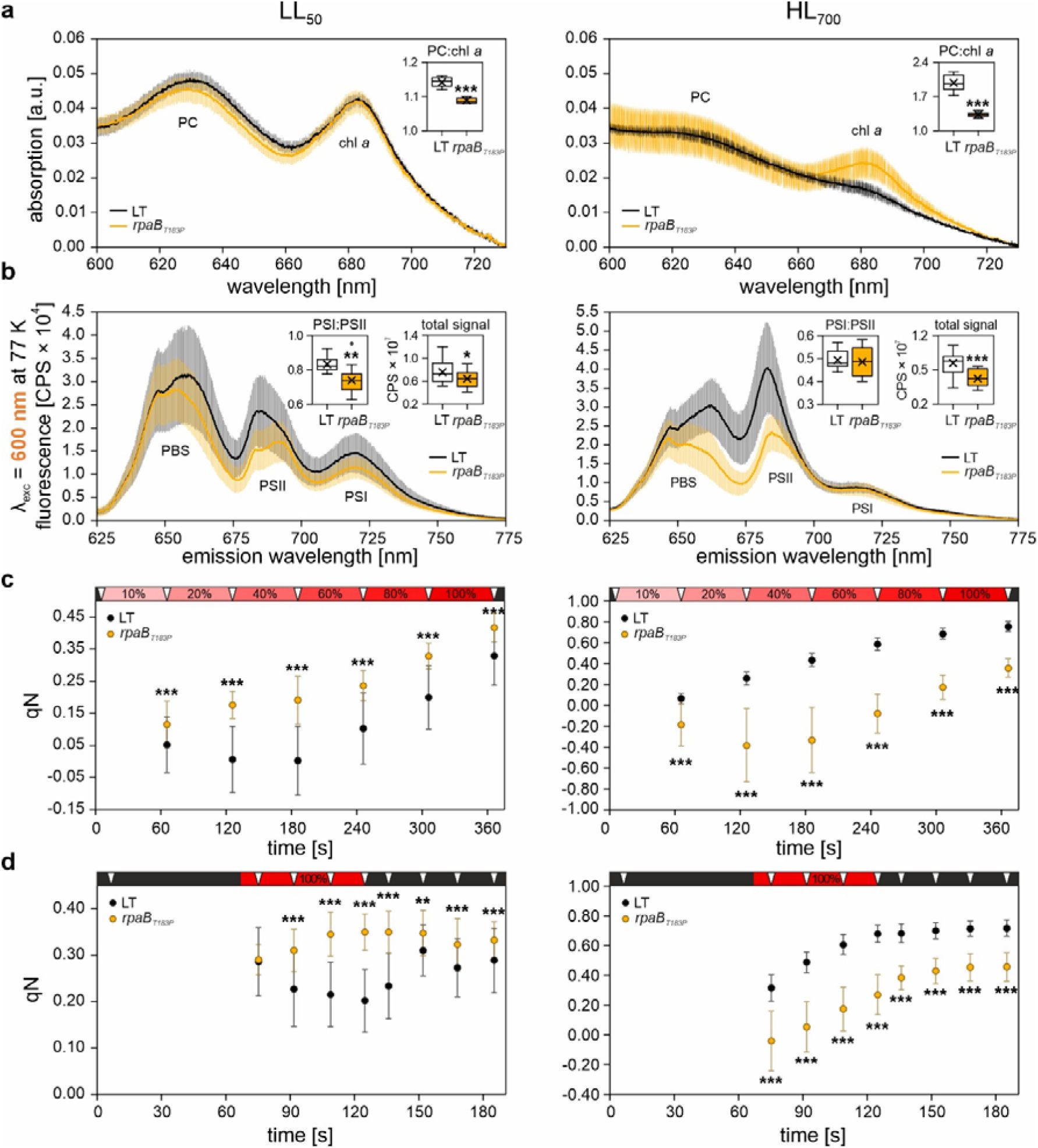
Spectroscopic characterization of *rpaB_T183P_*. **a** The absorption spectra of cultures grown under constant LL_50_ and HL_700_ in multi-cultivators, diluted to an optical density at 730 nm (OD_730nm_) of 0.1, were recorded at intervals of 0.2 nm). The absorption shoulders corresponding to phycocyanin (PC) and chlorophyll a (Chl a) are indicated. The inset box plot shows the distribution of PC:Chl a ratios, calculated from the corresponding absorption maxima as Abs_630nm_:Abs_684nm_ (LT and *rpaB_T183P_*under LL_50_, n = 4; LT and *rpaB_T183P_* under HL_700_, n = 12 and 24, respectively). **b** Fluorescence emission spectra were obtained by exciting undiluted samples (n = 18 biological replicates) with amber light (λ_exc_ = 600 nm, targeting peripheral antennae) at 77 K. Data were recorded at 0.25 nm intervals and are presented as counts per second (CPS), after subtraction of the baseline signal (average from the range F780-F800). The inset box plots show PSI:PSII ratios as estimated from F725:F695 (left) and the data distribution of the total signal, calculated as the sum of all intensities ranging from F626 to F780 (right). **c, d** Fluorescence at room temperature was measured from dark-acclimated, OD_730nm_-normalised LT and *rpaB_T183P_* droplets under incremental red-orange illumination (**c**) and an induction-relaxation regime (**d**). Dark periods are indicated in black, actinic red-orange light (100% ∼ 215 µmol photons m^-2^ s^-1^; λ_max_ = 625 nm) in red, and saturating pulses (cold-white 6500 K LEDs providing ∼1200 µmol photons m^-2^ s^-1^) as downward triangles in white. The coefficient of non-photochemical quenching of fluorescence (qN) was automatically calculated mid- and post-illumination after each saturating pulse (LT and *rpaB_T183P_*under LL_50_, n = 40; LT and *rpaB_T183P_* under HL_700_, n = 12 and 24, respectively). Statistical significance was determined using Student’s t-Test, with p ≤ 0.05 (*), p ≤ 0.01 (**), and p ≤ 0.001 (***).

PSI:PSII ratios were lower in LL_50_-acclimated *rpaB_T183P_*cells compared to LT but identical in HL_700_-acclimated cells (**Fig. 9b**), consistent with the D1 and PsaC immunoblot signal quantification (**Fig. 8b**). These observations suggest a differential effect of *rpaB_T183P_* on photosystem stoichiometry and peripheral antennae accumulation, with PSII accumulation promoted under continuous LL and phycobilisome accumulation reduced under LL and moderate HL.

Finally, in vivo fluorescence measurements revealed that non-photochemical quenching (qN)^47^ in *rpaB_T183P_* cells increased in LL_50_-acclimated cells and decreased in HL_700_-acclimated cells relative to LT (**Fig. 9c, d**). These results suggest that state transitions, the main contributor to qN in cyanobacteria^38,48^, were increased in LL_50_-acclimated and decreased in HL_700_-acclimated *rpaB_T183P_* mutants compared to the LT control.

## DISCUSSION

Fluctuations in light intensity represent one of the most rapid and severe environmental stressors that photosynthetic organisms must cope with, disrupting carbon assimilation and causing damage to photosystems^25,49^. Consequently, the molecular mechanisms underlying FL tolerance and their application in crop improvement are being intensively investigated. ALE using cyanobacteria as chloroplast proxies has been previously employed to address HL-related stress^34,35,36^. In this study, evolutionary screening under two complex FL regimes was applied to identify new FL tolerance factors and adaptive alleles. Among 412 candidate mutations, three non-synonymous SNPs in genes encoding the protein Sll0518 with unknown function, the PSII assembly factor Pam68, and the Response Regulator RpaB, were reconstituted in the parental LT background and confirmed to confer varying yet specific FL adaptation, thus demonstrating that FL tolerance can be improved through ALE.

The A133V mutation in Sll0518 was shown to promote growth under both non-lethal FL0 and LL_12_ conditions, but not HL_1200_ (see **Fig. 4** and **Supplementary** Fig. 2), indicating a specific role in FL acclimation. This mutation was observed in all monoclonal strains (see **Table 1**, **Fig. 3**), further supporting its adaptive nature. Sll0518 has recently been found to co-immunoprecipitate with the RNA recognition motif protein Rbp3 and is encoded directly downstream of the *rbp2* gene in the Synechocystis genome^50^. In the same study, Rbp3 was found to bind mRNAs encoding photosynthetic proteins and to localize near the ribosome, implying an involvement in the regulation of photosynthetic protein translation, with Δ*rbp3* mutants displaying a lowered PSI:PSII ratio^50^. Although the precise function of Sll0518 remains unclear, it may play a supportive role in the biogenesis of PSI, a key target of FL-induced photodamage^25^.

The S113G mutation in the PSII assembly factor Pam68^40^, present in all 24 monoclonal strains, also enhanced tolerance to FL0, but did not improve growth under LL_12_ or HL_1200_ conditions (see **Fig. 4** and **Supplementary** Fig. 2). Previous studies have reported a lethal phenotype in Pam68 deletion mutants exposed to alternating darkness and HL^51^, further indicating a role for Pam68 in FL tolerance. Overexpression of WT Pam68 also resulted in enhanced FL0 tolerance, although to a lesser extent than the Pam68_S113G_ mutant strain (see **Fig. 5**). While total Pam68_S113G_ levels were found depleted under FL0, thylakoid-associated levels were unaltered or slightly increased (see **Fig. 6**), suggesting that the mutant protein may have an enhanced affinity for its thylakoid protein targets and that overexpression of the WT Pam68 protein might improve FL tolerance through increasing total and thylakoid-associated Pam68 levels. The Pam68_S113G_ mutation also stimulated the accumulation of photosynthetic proteins (D1, PsaC, and the phycobiliproteins PC and APC) under LL_50_ conditions, see **Fig. 6**). As no improvement in growth under LL_12_ conditions could be observed (see **Supplementary** Fig. 2), carbon assimilation, rather than light energy harvesting and conversion, may be the limiting factor for growth under LL. This is consistent with previous reports that overexpression of phosphoenolpyruvate carboxylase enhances growth at very low light intensities^52^, and proteomic analyses indicating that carbon assimilation proteins are more responsive to changes in light intensity than to changes in inorganic carbon availability^16^.

Pam68 promotes the accumulation of the PSII assembly intermediates RCa and RCb^40^, CP47 biosynthesis and chlorophyll ligand insertion^51^. While the Pam68_S113G_ mutation did not affect the accumulation of the PSII assembly intermediate RC47, it did lead to a depletion of dimeric PSII under FL0 conditions (see **Fig. 6**). As dimeric PSII is more abundant in the Pam68 deletion mutant *ins0933* mutant^40^, this implies that the Pam68_S113G_ mutation has an opposite effect compared to Pam68 depletion. In line with this, the reduction in RCa assembly intermediates found in both *ins0933*^40^ and the *pam68_S113G_* mutant (see **Fig. 6**), can be explained by the Pam68 depletion destabilizing RCa, while Pam68_S113G_ accelerates its assembly into mature PSII, resulting in superficially similar effects on cellular RCa pools. Together with an apparent shift of the Pam68_S113G_ protein towards higher molecular weight fractions under FL0 (see **Fig. 6)**, this may indicate that the increased FL tolerance is due to an increased affinity of Pam68_S113G_ for its thylakoid-localized substrate. In fact, the Pam68_S113G_ mutation may lead to a stronger physical association of Pam68_S113G_ with nascent PSII, potentially masking the PSII monomer-monomer interface and resulting in decreased PSII dimerization. Given that Synechocystis PSII monomers have been reported to exhibit lower activity compared to PSII dimers^43^, the enhanced FL0 tolerance observed in the *pam68_S113G_*strain could partially arise from indirect PSI protection. This protection may occur through reduced PSII electron pressure due to an increased PSII monomer to dimer ratio under FL0 conditions in *pam68_S113G_*. This scenario bears resemblance to the situation in *Arabidopsis thaliana*, where numerous PSII-affecting suppressor mutations were identified in *pgr5* mutant suppressor screens under a lethal FL regime^53^. Nevertheless, the pronounced similarities between the growth effects of WT Pam68 overexpression and *pam68_S113G_*, despite lowered Pam68 levels in the latter, suggest that S113G is likely a gain-of-function mutation.

The RpaB protein, encoded by the Synechocystis open reading frame *slr0947*, plays a crucial role in regulating PSI gene expression. It binds to the high-light regulatory 1 (HLR1) sequence in PSI gene promoters, acting as a positive transcription regulator when phosphorylated under LL conditions and as a negative regulator when dephosphorylated under HL conditions^46,54,55,56^. RpaB is also redox-regulated through a thiol switch, with active dimers dissociating into presumably less active monomers upon reduction by thioredoxin, linking its activity to the redox state of the photosynthetic transport chain^54,56^.

We identified two non-synonymous mutations in the *rpaB* gene, one of which was evaluated for its adaptive value under FL conditions (**Table 1**). Notably, neither mutation affected the potential thiol switch residue (Cys59) or the likely site of RpaB phosphorylation (Ser198^57^). Mutant strains expressing RpaB_T183P_ remained viable under FL+ conditions, and grew comparable to the LT strain under LL_50_ conditions, while showing no difference in performance under FL0 conditions (see **Fig. 4**). The *rpaB_T183P_* strains were unique among reconstituted FL mutants in their ability to grow under HL_1200_ conditions, albeit with lower growth rate at LL_12_ conditions compared to LT (see **Supplementary** Fig. 2), suggesting a performance trade-off under LL while enhancing tolerance to HL in both continuous and fluctuating applications.

Interestingly, RpaB_T183P_ levels were significantly increased under both LL_50_ and HL_700_ conditions. Previous studies have shown that *rpaB* knock-down impairs growth under LL but promotes growth under HL conditions^46,58^. However, in our study, neither knock-down (*rpaBkd*) nor overexpression (*rpaBoe*) of *rpaB* had positive effects on growth under HL_1200_ and FL+ conditions (see **Fig. 7**), demonstrating that the single amino-acid exchange in RpaB confers additional functionality to RpaB_T183P_.

A possible mechanism for HL and FL+ tolerance induced by RpaB_T183P_ involves downregulation of photosynthetic proteins (D1, PsaC, and APC+PC) under elevated light intensities (see **Fig. 8**), suggesting an overall depletion of PSII, PSI, and phycobilisomes. This aligns with observations from suppressor screens in *Arabidopsis thaliana pgr5* mutants, where increased FL sensitivity was overcome through mutational disruption of the photosynthetic electron transport chain to prevent PSI damage^53^. However, unlike the findings in this study, downregulation of PSI did not increase FL tolerance in Arabidopsis^53^.

The *rpaB_T183P_* mutant strains exhibited a notable decrease in PC:Chl a ratios, particularly under HL_700_ conditions (see **Fig. 9**), suggesting that increased FL tolerance is linked to reduced light energy harvesting capacity through peripheral antennae. This was accompanied by a significant reduction in phycobilisome and PSII fluorescence emission at 77 K (see **Fig. 9**), consistent with previous findings in *rpaB* knockdown strains^41^. Under HL_700_ conditions, the mutant cells showed a marked decrease in non-photochemical quenching (qN)^59^ (see **Fig. 9**), likely due to reduced PSII and phycobilisome levels. In contrast, under LL_50_ conditions, the mutant displayed increased qN. Since state transition is considered to be the main contributor to NPQ in cyanobacteria^38,59^, this may indicate a reduced excitation transfer from phycobilisomes to PSII in LL_50_ acclimated cells, due to a greater transition from state 1 (where phycobilisomes primarily transfer excitation energy to PSII) to state 2 (where phycobilisomes primarily transfer excitation energy to PSI).

In the *rpaB_T183P_* mutant, PSI marker protein PsaC accumulation remained unaffected under LL_50_ conditions but showed a decrease under HL_700_ conditions (see **Fig. 8**). This suggests that the RpaB_T183P_ protein maintains its full functionality under LL conditions, while displaying increased activity in its HL-induced form, which represses PSI gene expression. The observed reduction in PSII and phycobilisome levels under HL_700_ conditions is unlikely to be a direct consequence of the decrease in PSI levels. Previous research has shown that PSII levels do not closely track changes in PSI levels, whereas PSI levels tend to follow alterations in PSII levels more closely^60,61^. This observation, combined with improved viability of the mutant under HL and FL conditions, suggests that the RpaB_T183P_ substitution is likely a gain-of-function mutation. Considering that RpaB-like proteins are conserved across cyanobacterial and plastid phylogeny, their homologs in eukaryotic algae and higher plants (known as Ycf27 proteins^44^) could serve as potential targets for enhancing FL tolerance in crop plants.

In conclusion, our findings demonstrate that each of the three candidate mutations examined in this investigation resulted in improved FL tolerance when introduced into the original Synechocystis LT strain. This outcome underscores the effectiveness of ALE as a method for identifying new genetic variants that confer increased resilience of photosynthetic processes in the face of abiotic stresses. The pioneering application of fluctuating light intensities in this study opens up possibilities for developing even more intricate stress regimes in future research. It is particularly noteworthy that this approach successfully targeted both genes with unknown functions that are unique to cyanobacteria, as well as genes encoding proteins that have homologs in the chloroplasts of flowering plants. This dual targeting allows for the assignment of functions to previously uncharacterized proteins and facilitates the discovery of genetic variants that could potentially be applied in the engineering of photosynthetic processes in crop plants.

## METHODS

### Synechocystis strains: generation and culture conditions

*Synechocystis* sp. PCC 6803 glucose-tolerant cells, referred to as “laboratory type” (LT) were kindly provided by Himadri Pakrasi (Washington University, St. Louis, USA). Previously described knock-out and knock-down mutants of *pam68* and *rpaB47* were utilized^40,46^. Reconstruction of *sll0518_A133V_*, *pam68_S113G_*, *rpaB_T183P_* in the LT background employed established methodology^35^ with plasmid vectors constructed using a pUC57-mini vector backbone derived from IMBB2.4CpUC57Cmini kindly provided by Professor Neil Hunter (University of Sheffield). Overexpression mutants of *pam68* and *rpaB* were generated through homologous recombination using non-replicative vectors derived from IMBB2.4-pUC57-mini and pICH69822 (obtained from E. Weber, Icon Genetics GmbH, Halle, Germany), respectively. These constructs were assembled via Gibson assembly and targeted to the genomic neutral site *slr0168*, with *pam68* and *rpaB* coding sequences expressed under the control of the strong *psbA2* and *rbcL* promoters, respectively.

Cultures were typically grown under continuous illumination at 30 μmol photons mC² sC¹ of white fluorescent light (OSRAM HE28W/830 Lumilux warm white Hg fluorescent lamps) at 23 °C. Liquid cultures were inoculated at an initial OD = 0.05 in BG11 photoautotrophic medium, with 5 mM glucose added for pre-transformation cultures. Growth was conducted in Multi-Cultivator MC 1000-OD devices, equipped with an AC-700 cooling unit and a warm-white LED panel (Photon System Instruments, Drasov, Czech Republic). For solid media growth, BG11 was supplemented with 0.75% (w/v) bacteriological agar.

### Adaptive evolution of Synechocystis under fluctuating light

Two FL adaptive evolution experiments were conducted using Synechocystis, relying on its natural mutation rates^35^ to identify new adaptive alleles. The experiments began with six separate batch cultures derived from a Synechocystis LT stock culture. Each culture was grown in a 100 mL glass tube within a temperature-controlled water bath of a multicultivator. The propagation cycles involved 70 mL of medium with an initial OD_730nm_ of 0.05, incubated under constant aeration at 23 °C with fluctuating warm-white LED illumination for 7-14 days. The light fluctuations were progressively intensified in both amplitude and frequency throughout the selection process.

The FL0 regime alternated between 1 min of high light (HL) and 5 min of low light (LL) throughout the entire adaptive laboratory evolution (ALE) protocol. The selection process began with five cycles of 50 µmol photons m^-2^ s^-1^ (LL_50_) and 700 µmol photons m^-2^ s^-1^ (HL_700_), followed by three cycles with varying LL and HL intensities. The final 12 cycles used a regime of 5 min at LL_12_ and 1 min at HL_1200_.

The FL+ regime also started with 1 min of HL followed by LL, but the LL period was progressively shortened from 5 min to 1 min. The initial cycle matched that of the FL0 regime, with subsequent cycles gradually reducing LL intensity and increasing HL intensity. The final 12 cycles alternated between 1 min of LL_12_ and 1 min of HL_1200_.

Both FL0 and FL+ experiments consisted of 20 selective cycles in total, resulting in three evolved batch cultures for each condition, labeled FL0_a_20_, FL0_b_20,_ FL0_c_20_ and FL+_a_20_, FL+_b_20_, FL+_c_20_, respectively (**Fig. 1a**). Further details of the selective cycle protocols can be found in **Supplementary Table 1**.

### Isolation of FL-adapted Synechocystis clones for genome re-sequencing

Single clones were isolated by plating dilutions (10^-6^-10^-7^) of FL0_a_20_, FL0_b_20_, FL0_c_20_ and FL+_α_20_, FL+_β_20_, FL+_γ_20_, onto solid BG11 media. Isolation plates were incubated at 30 µmol photons m^-2^ s^-1^ continuous illumination and 23 °C for seven days. Representatives of mutant subpopulations were sampled by selecting different clones based on colony color and size and isolated clones were grown on solid BG11 media for seven days.

Subsequently, clones for genome re-sequencing were selected as previously described^35^ based on room-temperature fluorescence parameters measured by FluorCam 800MF (Photon Systems Instruments, Drasov, Czech Republic). For FL0 and FL+, *n* = 66 and *n* = 72 isolated clones were assessed. To capture the genetic variability within each batch culture, clones best representing the quartiles of the observed the Fv^-^/Fm^-^ distributions (*i.e.*, two extreme and two intermediate values of PSII quenched quantum efficiency) were selected for whole-genome resequencing, totaling *n* = 12 for FL0 and FL+, respectively.

### Synechocystis genomic DNA extraction and sequencing

Genomic DNA for wholeCgenome sequencing was isolated from cell pellets of 10–30 mg fresh weight following the manufacturer’s protocol (EasyPure® Plant Genomic DNA Kit, TransGen Biotech Co., Ltd., Beijing, China). Cells were broken in EasyPure®lysis buffer using a 1:1 mixture of small glass beads (425–600 µm + 212–300 µm, Sigma Aldrich, St. Louis, MO, United States) and a TissueLyser II (QIAGEN, Hilden, Germany). DNA isolates were then subjected to agarose gel electrophoresis to assess structural integrity. The genomes of 24 monoclonal mutants (four per adapted batch culture) were then reCsequenced on the Illumina HiSeq platform (2 × 150Cbp pairedCend reads) by NovoGene Ltd. (Cambridge, United Kingdom).

### Sequence data quality control and filtering

The previously published Synechocystis LT_t=0_ genome assembly^35^ served as the control for excluding background mutations; all mutations identified in FL-ALE were tracked relative to the LT_t=0_ assembly.

Adapting previously described methodology^35^, the quality of the WGS raw data was assessed using FastQC v0.11.9^62^. Pre-processing began with Cutadapt v4.1^63^, which filtered out low-quality reads and removed sequences containing adapter contamination or more than 10% undetermined bases (‘N’ bases). Rcorrector^64^ was then used to perform *k*-mer correction on the filtered datasets, applying the default *k*-mer length setting. The resulting dataset, consisting of filtered and corrected reads, was used for subsequent mutation detection. Genome resequencing of the 24 single clones yielded an average coverage of 339±44/638±137/300±65/116±42/577±193-fold for chromosome/pSYSM/pSYSA/pSYSG/pSYX, respectively.

### Variant analysis

The Breseq pipeline^65^ were applied for the identification of potential mutations. The clean reads were aligned to the *Synechocystis* sp. PCC 6803 reference genome (ASM972v1) obtained from the NCBI database using bowtie2 v2.5.1^66^. The generated SAM alignment files were then used for variant calling. Breseq analysis was conducted in two distinct modes. The ‘consensus’ mode defined a mutation as ‘fixed’ when its frequency was ≥0.80, while considering a site as ‘polymorphic’ when the variant frequency ranged between 0.20 and 0.80. On the other hand, in the ‘polymorphism’ mode, a mutation was designated as ‘fixed’ at frequencies ≥0.95 and as ‘polymorphic’ if its occurrence spanned frequencies between 0.05 and 0.95. For an overview of fully segregated, protein-affecting mutations identified in FL-ALE strains, see **Table 1**.

### Phylogenetic analysis

The phylogenetic analysis was carried out considering 101 polymorphic sites representing all deviations from the reference genome (ASM972v1) with a 100% frequency (*i.e*., identified as fully segregated in at least one sample). IQ-TREE multicore version 2.2.6^67^ was used to perform the subsequent phylogenetic analysis. Briefly, the model selection method was applied with default parameters to identify the most suitable model for the data set. Afterwards, the selected model (according to Bayesian Information Criterion “BIC” values), that is Kimura 2 Parameter (K2P) with equal frequencies, was applied to infer the maximum likelihood phylogenetic relationships between the samples. The bootstrapping method was applied to validate the generated tree with 500 replicates. The resulting phylogenetic tree was generated using CLC Main Workbench (QIAGEN, Venlo, Netherlands).

### Pigment extraction and quantification, determination of phycocyanin:chlorophyll ratios

Chlorophyll *a* (Chl *a*) and total carotenoids (Cars)), were extracted and quantified as previously described^35^. Molar ratios of the peripheral antenna pigment phycocyanin (PC) to core antenna pigment chlorophyll *a* in cultures seven days past inoculation were estimated as previously described^35^.

### Determination of PSII quantum yield parameters

Apparent PSII quantum yield was measured as Fv^-^/Fm^-^^38^ at room temperature as previously described^35^ using a FluorCam 800MF (Photon Systems Instruments, Drásov, Czech Republic). Fv-/Fm- was measured in cells grown for 7 days at 23 °C and 30 µmol photons m^-2^ s^-1^ on BG11 solid media after single-clone isolation, or cells collected from cultures grown at 50 µmol photons m^D2^ s^D1^ in MC and subsequently normalized to OD_730nm_ = 10 of which 10 µL droplets were placed on BG11-agar. After overnight acclimation at 30 µmol photons m^-2^ s^-1^ and 23 °C, plates were dark-incubated for one hour before measurements. Instrument sensitivity was manually adjusted (20%–30%) before measuring. For samples under HL, suspensions were collected from the liquid cultures after 7 day cultivation and prepared as described previously^8^.

### Protein extraction, detection and quantification

Cells were collected from a 1-mL suspension at OD_730nm_ = 10 by gentle centrifugation, and the pellets were snap-frozen in liquid N_2_ and stored at −80 °C. The cell pellets were then homogenized and lysed twice in 500 µL of homogenization buffer (0.4 M sucrose, 10 mM NaCl, 5 mM MgCl_2_, 20 mM Tricine, adjusted to pH 7.9 with HCl), supplemented with protease inhibitors (cOmplete™ Mini EDTA-free Protease Inhibitor Cocktail, Roche AG, Basel, Switzerland) and approximately 1 mL of a glass bead mixture. Lysis was performed using a mixer mill (MM 400, RETSCH, Haan, Germany) with five cold-lysis cycles (5 min at 30 Hz). After centrifugation at 4 °C, the supernatant was collected, and protein concentration was estimated using Bradford (ROTI-Quant, Carl Roth, Karlsruhe, Germany) and BCA (Pierce™ BCA Protein Assay Kit, Thermo Fisher Scientific, Waltham, MA, USA) protein assays. Samples were stored at −20 °C until further processing.

For SDS-PAGE, protein extracts (20 µg of protein) were mixed with 5× SDS loading dye, denatured (65 °C, 10 min) and size-separated on 10% Tris-Tricine gels. Phycocyanin and allophycocyanin were quantified by recording fluorescence (λ_emission_ ≥ 600 nm) directly from the gels^68^ (Fusion FX imaging system, Vilber, Collégien, France; excitation wavelength of λ_excitation_ = 530 nm). Image analysis and signal quantification were conducted using ImageJ^69^. Proteins were then transferred to PVDF membranes (Immobilon-PSQ, Millipore, Burlington, MA, USA) via electroblotting. Membranes were stained with Coomassie Brilliant Blue (CBB) for loading control, then de-stained before immunodetection.

For specific protein detection, membranes were cut or used whole, blocked with 1.5% (w/v) BSA in TBST, and incubated with primary antibodies against PsaC and PsbA (Agrisera, Vännäs, Sweden), Pam68 (kindly provided by Prof. Dr. Jörg Nickelsen, LMU Munich, Germany), and RpaB (PhytoAB, San Jose, CA, USA). After overnight incubation with primary antibodies and 2-hour incubation at 4 °C with horseradish-peroxidase coupled secondary antibodies, chemiluminescence was detected using SuperSignal™ West Pico PLUS chemiluminescent substrate (Thermo Fisher Scientific, Waltham, MA, USA) and imaging system (Fusion FX imaging system, Vilber, Collégien, France). Signal quantification was performed using ImageJ software^69^.

### Size separation of thylakoid protein complexes by Blue Native PAGE and Second-Dimension SDS PAGE

Thylakoid fractions and native page gels were prepared as previously described^42^. Sample aliquots containing approximately 5 µg of chlorophyll were supplemented with 1% β-DM, and non-solubilized material was removed by centrifugation. The supernatant was subjected to Blue Native PAGE separation (first dimension) using 1 mm gels with a 4–12% (v/v) acrylamide gradient (40% w/v acrylamide) in gel buffer (3 M aminocaproic acid, 300 mM Bis-Tris, adjusted to pH 7.0 with HCl). Electrophoresis was performed at a constant voltage of 100 V using a cathode buffer (50 mM Tricine, 15 mM Bis-Tris, 0.02% CBB G-250, adjusted to pH 7.0 with HCl) and an anode buffer (50 mM Bis-Tris, pH 7.0). After electrophoresis, gel lanes were excised and stored at C20 °C until further processing.

For size-separation of denatured thylakoid complex components by second-dimension SDS PAGE, gel lanes were incubated in fresh Tris-HCl buffer (25 mM Tris, 1% SDS, 1% DTT, pH 7.5) for 20 min under gentle shaking. The samples were then subjected to SDS-PAGE using 10% Tris-Tricine SDS polyacrylamide gels at a constant voltage of 55–85 V. The resulting gels were processed for protein transfer via semi-dry electroblotting and immunodetection, following the procedure detailed above.

### Low-temperature fluorescence spectrometry

Low-temperature fluorescence spectra of *Chl a* and phycobiliproteins were recorded using a HORIBA Fluoromax Plus FL-1013 spectrofluorometer (HORIBA Jobin Yvon GmbH, Oberursel, Germany). Cultures grown in MC photobioreactors under the specified light conditions were transferred into glass capillaries (Hilgenberg GmbH, Malsfeld, Germany) directly from the cultivation device at 7 days past inoculation and immediately snap-frozen in liquid nitrogen. Samples were stored in a light-occluding casing at −80 °C for 1–2 weeks and measured in a single batch. Measurements were conducted at 77 K in a dewar filled with liquid nitrogen, using a signal integration time of 0.2 s nmC¹ and a detection bandwidth of 1 nm. Fluorescence signals were detected through a 500 nm long-pass filter upon excitation at 435 nm (λ*exc* = 435 nm) to stimulate Chl *a* absorption and at 600 nm (λ*exc* = 600 nm) to selectively excite phycobiliproteins.

Key fluorescence peaks were analyzed as ratios derived from baseline-subtracted spectra, where the baseline was defined as the mean fluorescence (F) between 780 and 800 nm emission wavelength (F780, F800). Relevant peaks corresponded to phycocyanin (PC; F645), allophycocyanin (APC; F662), Chl *a* in PSII core antenna CP43 (CP43; F685), *Chl a* in PSII core antenna CP47 (CP47; F695), and Chl *a* in PSI (PSI; F725) as described ^35^.

### Determination of coefficient of non-photochemical quenching at room temperature

To determine the coefficient of non-photochemical quenching at room temperature, fast fluorescence kinetics were measured using a FluorCam 800MF (Photon Systems Instruments, Drásov, Czech Republic). Cells collected from cultures grown at 50 µmol photons m^D2^ s^D1^ in MC were normalized to OD_730nm_ = 10, and 10 µL droplets were evenly distributed on BG11-agar. After overnight acclimation at 30 µmol photons m^-2^ s^-1^ and 23 °C, plates were dark-incubated for one hour before measurements. Instrument sensitivity was manually adjusted (20%–30%) before measuring fluorescence kinetics under incremental red-orange illumination (Fo 5 s > Fm pulse 0.8 s > AL 10 % 60 s > Fm pulse 0.8 s > AL 20 % 60 s > Fm pulse 0.8 s > AL 40 % 60 s > Fm pulse 0.8 s > AL 60 % 60 s > Fm pulse 0.8 s > AL 80 % 60 s > Fm pulse 0.8 s > AL 100 % 60 s > Fm pulse 0.8 s > END) and induction-relaxation regime (Fo 5 s > Fm pulse 0.8 s > dark 10 s > AL 100 % 60 s > dark relaxation 60 s; 9 Fm pulses during AL and dark relaxation, first Fm pulse 9 s after AL onset).

Actinic light at 100% intensity approximated 215 µmol photons m□² s□¹ (λ_max_ = 625 nm), while saturating pulses (Fm pulses; ∼1200 µmol photons m□² s□¹) were provided by cold-white 6500 K LEDs. Quenching parameters were computed using FluorCam7 (Photon Systems Instruments).

### Statistics and boxplot description

Statistical analyses were conducted using two-tailed Student’s *t*-tests in Microsoft Excel. Post-hoc Bonferroni-Holm corrections for multiple comparisons were performed using astatsa when significant differences among groups were detected via one-way ANOVA at https://astatsa.com/.

Box plots depicting data point distributions were created in Microsoft Excel. The horizontal middle lines represent inclusive medians, crosses indicate mean values, boxes denote the second and third quartiles, whiskers extend to the first and fourth quartiles, and points beyond the whisker ranges indicate outliers exceeding 1.5 times the interquartile range.

### Data availability

DNA-Seq data are available from the NCBI SRA database under xxxx (for review purposes see: https://dataview.ncbi.nlm.nih.gov/object/PRJNA1228058?reviewer=okcif0og6oj4lmdrap39qhi6uh)

## ACKNOWLEDGEMENTS

We thank Prof. Jörg Nickelsen (LMU Munich, Germany) for providing the Pam68 antibody and Prof. Yukako Hihara (Saitama University, Japan) for sharing the RpaB knock-down mutant (*rpaBkd*).

## AUTHOR CONTRIBUTIONS

D.L. and T.F.-G. conceived the project. D.L. provided the funding. D.L. and T.F.-G. designed all experiments, with support from M.D. T.F.-G. performed the experiments with support from W.C. (RpaB mutant generation and characterization) and M.Z. (Pam68 mutant characterization). E.M.A.-S. performed the DNA sequence analysis and interpretation of the results. D.L. wrote the manuscript with the support of M.D. and T.F.-G. All authors read and approved the final manuscript.

## ADDITIONAL INFORMATION

**Supplementary information** The online version contains supplementary material available at xxxx.

