## Supplementary for "Improving tolerance to fluctuating light through adaptive laboratory evolution in the cyanobacterium Synechocystis"

**Supplementary Tables**

**Supplementary Table 1** The FL0 and FL+ selective regimes for FL-ALE.

The light regimes implemented during the 20 propagation rounds of the two FL-ALE protocols (FL0 and FL+) are detailed. The parameters include: LL/HL, low/high light intensities, quantified in μmol photons m^−2^ s^−1^; A, Amplitude, calculated as HL - LL; t_HL/LL_, duration of exposure to LL/HL expressed in min; and Cycles, number of propagation rounds. Throughout all experimental protocols, environmental conditions were standardized with an aeration rate of 100–150 mL of air per minute and a constant temperature of 23°C.

| **Condition** | **LL** | **HL** | **A** | **t_LL_** | **t_HL_** | **Cycles** |
| --- | --- | --- | --- | --- | --- | --- |
| **FL0** | | | | | | |
| initial | 50 | 700 | 650 | 5 | 1 | 5 |
| intermediate | 50 | 1000 | 950 | 5 | 1 | 1 |
|  | 50 | 1200 | 1150 | 5 | 1 | 1 |
|  | 20 | 1200 | 1180 | 5 | 1 | 1 |
| final | 12 | 1200 | 1188 | 5 | 1 | 12 |
| **FL+** | | | | | | |
| initial | 50 | 700 | 650 | 5 | 1 | 1 |
| intermediate | 50 | 700 | 650 | 4 | 1 | 1 |
|  | 50 | 700 | 650 | 3 | 1 | 1 |
|  | 50 | 700 | 650 | 2 | 1 | 1 |
|  | 50 | 700 | 650 | 1 | 1 | 1 |
|  | 50 | 1000 | 950 | 1 | 1 | 1 |
|  | 50 | 1200 | 1150 | 1 | 1 | 1 |
|  | 20 | 1200 | 1180 | 1 | 1 | 1 |
| final | 12 | 1200 | 1188 | 1 | 1 | 12 |

**Supplementary Figures**

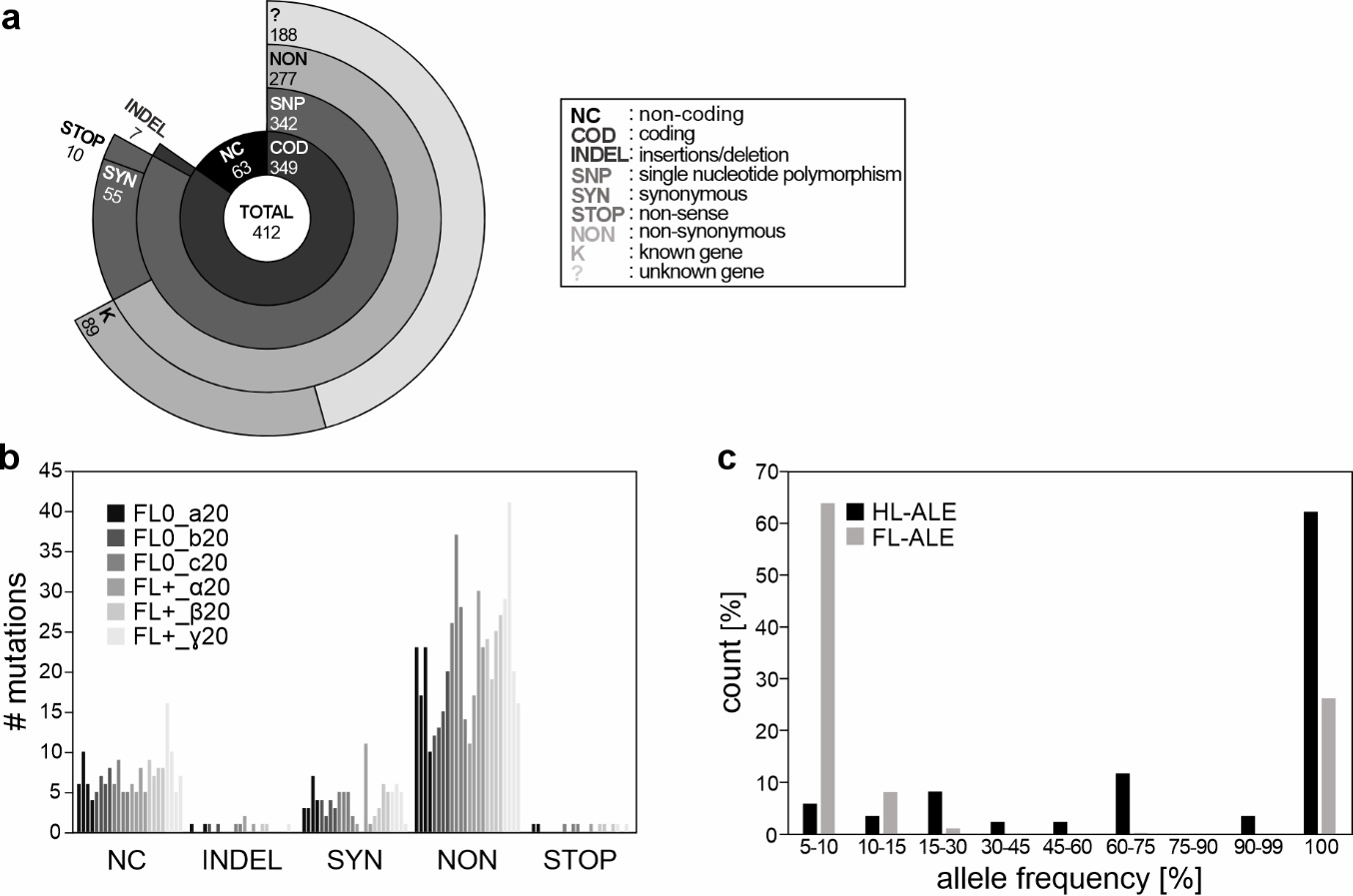

**Supplementary Fig. 1** **l** **Analysis of the mutational landscape observed in FL-adapted strains.**

**a** A hierarchical sunburst plot illustrates the distribution of novel alleles (absent in LT and WT strains) across four levels of classification: 1. Genomic context: Coding (COD) vs. Non-coding (NC) regions. 2. Mutation type in coding regions: Single-nucleotide polymorphisms (SNP) vs. Insertion/Deletions (INDELs). 3. SNP effect: Non-synonymous (NON), Synonymous (SYN) or Premature stop codon (STOP). 4. Functional annotation of non-synonymous mutations: Known/characterised (K) vs. Unknown/uncharacterised (?) genes.

**b** Bar chart quantifying the relative abundance of mutations across five categories: NC, INDEL, NON, SYN, and STOP.

**c** The frequency distribution of the total mutations observed in the two ALE experiments is presented, categorized into nine bins. This distribution is compared to previously reported data from high light (HL) ALE experiments conducted without external mutagens (see main text).

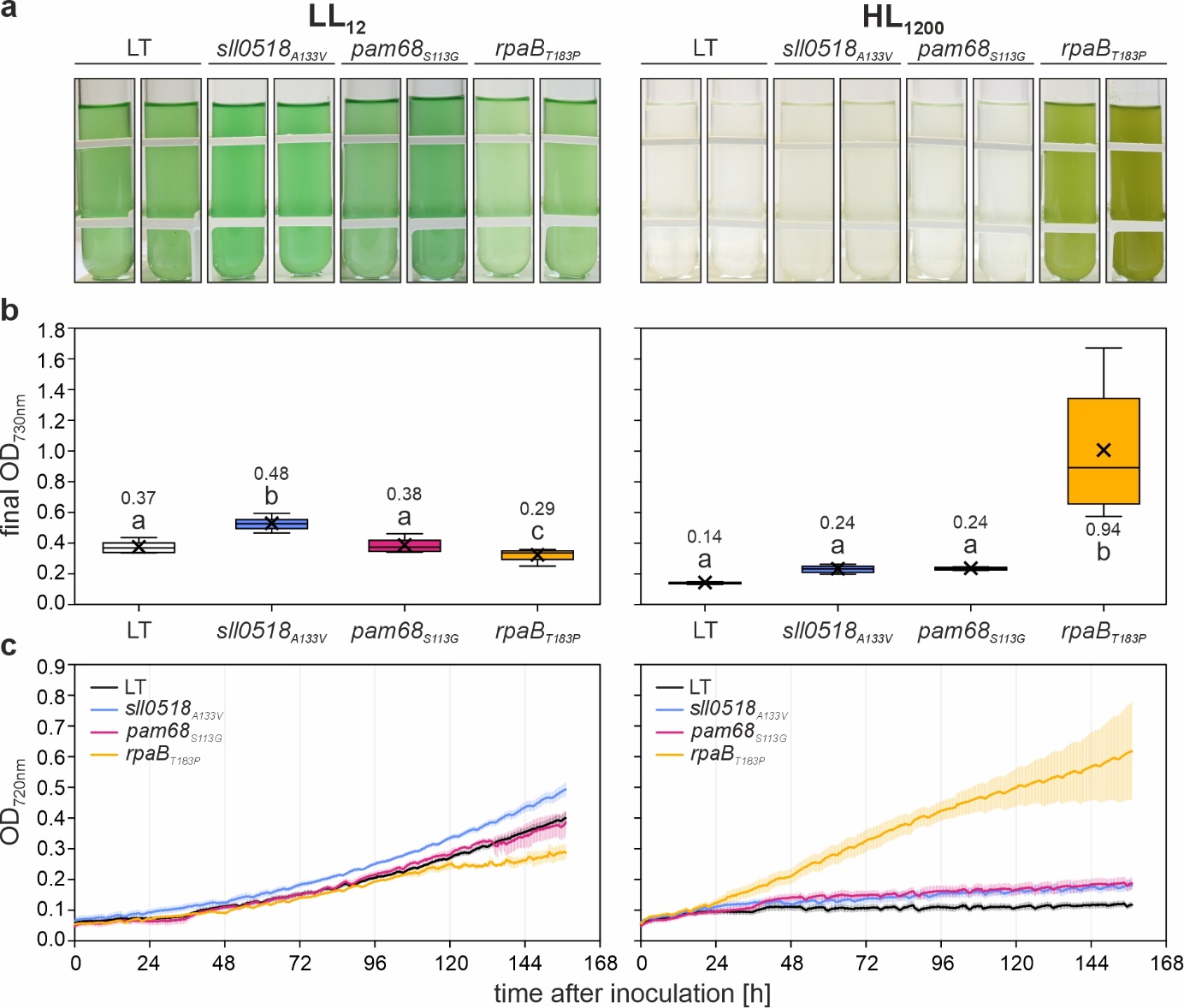

**Supplementary Fig. 2 l** **Growth characteristics of *sll0518_A133V_*, *pam68_S113G_*, *rpaB_T183P_* and LT strains under continuous LL_12_** **and HL_1200_ conditions.**

**a** Visual representation of liquid cultures for the four strains cultivated under LL (left) or HL (right) conditions at 23 °C with 100 mL min^-1^ aeration, photographed seven days after post-inoculation. Data were collected from cultures grown in multi-cultivators.

**b** Quantitative analysis of cell density (OD_730nm_) for the four strains grown under conditions described in A. Box plots are presented, with lowercase letters denoting statistically significant differences (p ≤ 0.05) as determined by post-hoc Bonferroni-Holm simultaneous comparison of all measurements (*n* = 8) following significant between-group differences detected by one-factorial ANOVA.

**c** Growth kinetics of the four strains under LL and HL conditions, monitored automatically by the multi-cultivators measuring OD_720nm_. Error bars represent the standard deviation (n = 4).

Statistical data in panel b are presented as box plots, showing individual data points, median (horizontal lines), mean (crosses), interquartile range (box), and 1.5× interquartile range (whiskers).

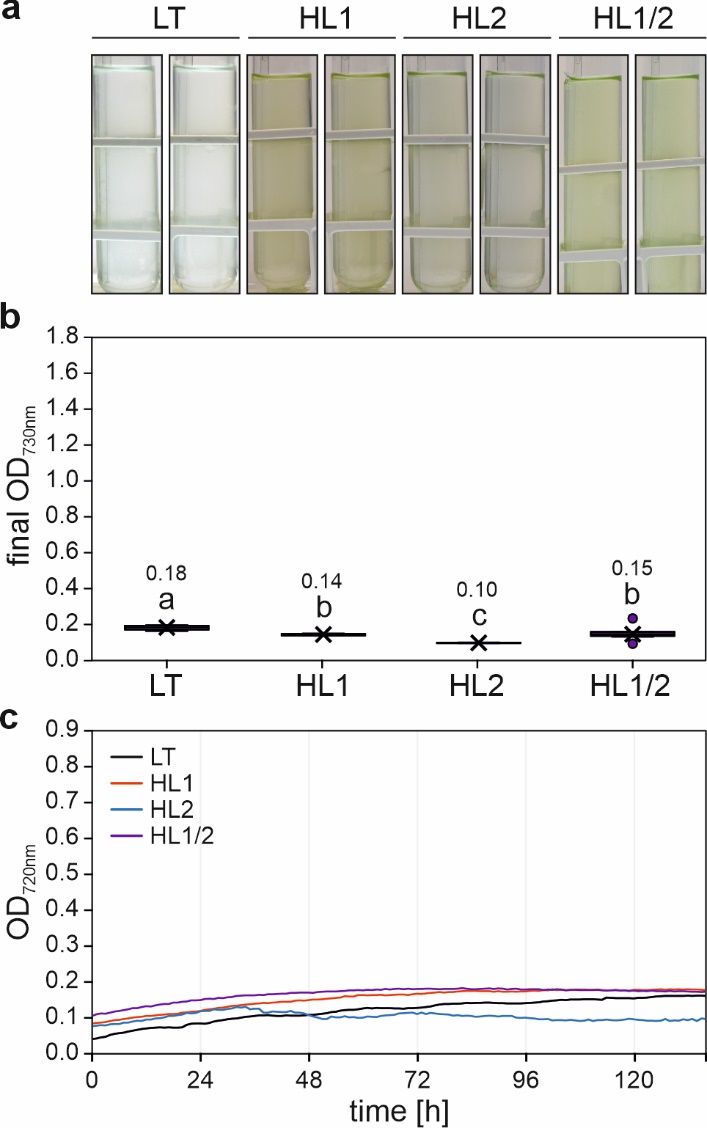

**Supplementary Fig. 3** **l** **Growth characteristics of HL-tolerant strains HL1, HL2, HL1/2 and LT strains under final FL+ conditions.**

**a** Visual representation of liquid cultures for the four strains cultivated in multi-cultivators under final FL+ conditions at 23 °C with 100 mL min^-1^ aeration, photographed seven days post-inoculation.

**b** Quantitative analysis of cell density (OD_730nm_) for the four strains grown under conditions as in A. Box plots are presented, with lowercase letters denoting statistically significant differences (p ≤ 0.05) as determined by post-hoc Bonferroni-Holm simultaneous comparison of all measurements (n = 8) following significant between-group differences detected by one-factorial ANOVA.

**c** Growth kinetics of the four strains under final FL+ conditions, monitored automatically by multi-cultivators measuring OD_720nm_. Error bars represent the standard deviation (*n* = 4).

Statistical data in panel b are presented as box plots, showing individual data points, median (horizontal lines), mean (crosses), interquartile range (box), and 1.5× interquartile range (whiskers).
